# scReQTL: an approach to correlate SNVs to gene expression from individual scRNA-seq datasets

**DOI:** 10.1101/2020.07.13.200956

**Authors:** Hongyu Liu, N M Prashant, Liam F. Spurr, Pavlos Bousounis, Nawaf Alomran, Helen Ibeawuchi, Justin Sein, Piotr Słowiński, Krasimira Tsaneva-Atanasova, Anelia Horvath

**Affiliations:** McCormick Genomics and Proteomics Center, School of Medicine and Health Sciences, The George Washington University, 20037 Washington, DC, USA; Chinese Medicine Toxicological Laboratory, Institute of Traditional Chinese Medicine, Heilongjiang University of Chinese Medicine, Harbin 150040, PR China; Department of Medical Oncology, Dana-Farber Cancer Institute, Boston, MA 02215, USA; Cancer Program, The Broad Institute of MIT and Harvard, Cambridge, MA 02142, USA; Translational Research Exchange at Exeter, University of Exeter, Exeter, EX4 4QJ, UK; EPSRC Centre for Predictive Modelling in Healthcare, University of Exeter, Exeter, EX4 4QJ, UK; Department of Mathematics & Living Systems Institute, University of Exeter, Stocker Road, Exeter, EX4 4QD, UK; Dept. of Bioinformatics and Mathematical Modelling, Institute of Biophysics and Biomedical Engineering, Bulgarian Academy of Sciences, 105 Acad. G. Bonchev Str., 1113 Sofia, Bulgaria; Department of Biochemistry and Molecular Medicine, Department of Biostatistics and Bioinformatics School of Medicine and Health Sciences, George Washington University, 20037 Washington, DC, USA

**Keywords:** eQTL, ReQTL, scReQTL, single cell, VAF_RNA_, scVAF_RNA_, scRNA-seq, SNV, genetic variation, RNA-seq, single cell RNA sequencing, single cell RNA-seq

## Abstract

Recently, pioneering eQTLs studies on single cell RNA-seq (scRNA-seq) data have revealed new and cell-specific regulatory SNVs. Because eQTLs correlate genotypes and gene expression across multiple individuals, they are confined to SNVs with sufficient population frequency. Here, we present an alternative sc-eQTL approach – scReQTL - wherein we substitute the genotypes with expressed Variant Allele Fraction (VAF_RNA_) at heterozygous SNV sites. Our approach employs the advantage that, when estimated from multiple cells, VAF_RNA_ can be used to assess effects of rare SNVs in a single individual. ScReQTLs are enriched in known genetic interactions, therefore can be used to identify novel regulatory SNVs.

## Introduction

In recent years, single cell RNA-seq (scRNA-seq) has become an increasingly accessible platform for genomic studies (1). By enabling cell-level analyses, scRNA-seq has major advantages for studying gene-regulatory relationships. Among others, the ability to distinguish cell populations and to assess cell-type specific transcriptome features, have shown great potential to identify new regulatory networks (2–4). Furthermore, scRNA-seq enables the assessment of intracellular molecular relationships, which can reveal cell-specific gene-gene interactions and co-regulated genetic features (2,5,6). These relationships can be reflected in mutually correlated molecular traits, including gene expression (GE) and expression of genetic variants, such as Single Nucleotide Variants (SNVs).

A popular method to study SNVs effects on GE is eQTL (Expressed Quantitative Trait Loci), which is based on testing for a correlation between the number of alleles bearing the variant nucleotide at the position of interest, and the level of local (cis) or distant (trans) GE (7). eQTLs have been mapped by large-scale efforts such as Genotype-tissue Expression Consortium (GTEx), PsychENCODE, ImmVar BLUEPRINT, and CAGE, which have been instrumental in identifying SNVs affecting GE (8–12).

Recently, pioneering eQTL studies on scRNA-seq data have emerged. By utilizing the advantages of the single cell resolution, these studies have revealed many new regulatory SNVs, including those with cell-specific or transient effects (2–4,13–16). To assess GE, these methods employ approaches specific to single cell transcriptomics, including accounting for drop-outs, classification of cells by type, and assessments of progressive cell stages (2–4,13–16). SNV information is traditionally obtained from the genotypes across multiple individuals and encoded as the number of alleles (0, 1 or 2) bearing the variant nucleotide. Accordingly, eQTL analyses are confined to SNVs present in a sufficient number of individuals in the studied group, and frequently exclude variants with low minor allele frequency in the population.

Here, we explore an alternative approach to assess effects of SNVs on GE from scRNA-seq data, wherein we substitute the genotype counts with the proportion of expressed variant-bearing RNA molecules (Variant Allele Fraction, VAF_RNA_) at heterozygous SNV loci. Our approach employs the advantage that, when estimated from multiple cells, VAF_RNA_ can be used to assess effects of rare SNVs in a single sample or individual.

To correlate VAF_RNA_ to GE from single cells, we first identify heterozygous SNVs from the pooled RNA-sequencing data, then estimate VAF_RNA_ in the individual cell alignments, and correlate VAF_RNA_ with GE from the individual cells using a linear regression model (17). To develop the pipeline, we used recent methodologies for calling SNVs and VAF_RNA_ estimation from RNA-seq data (18–22), as well as scRNA-seq-specific methods to estimate GE (23). We also adopted a strategy from a method recently developed in our lab to correlate VAF_RNA_ and GE from bulk RNA-sequencing data – ReQTL (RNA-eQTL) (24). We term the application of this technique on single-cell RNA-sequencing data: scReQTL.

We applied scReQTL on publicly available scRNA-seq generated on the 10×Genomics Chromium platform using 3’-based protocol on 26,640 cells obtained from three healthy female donors (25). scReQTL analysis was performed after classification of the cells by cell type, and only SNVs covered by a minimum of 10 unique sequencing reads per cell were included in the analysis. Across the three samples, we identified 1272 unique scReQTLs. scReQTLs common between individuals or cell types were consistent in terms of the directionality of the relationship and the effect size. In addition, scReQTLs were substantially enriched in known gene-gene interactions and significant genome-wide association studies (GWAS) loci.

## Results

### Overview of scReQTL workflow

The overall scReQTL workflow is presented in Figure 1 and outlined in detail in the Materials and Methods section. The pipeline includes 5 major steps: scRNA-seq data processing, GE estimation, cell type identification, VAF_RNA_ assessment, and SNV-GE correlation by cell type. Below, we briefly describe elements that we identified as specific and essential for the scReQTL analysis.

**Figure 1.**
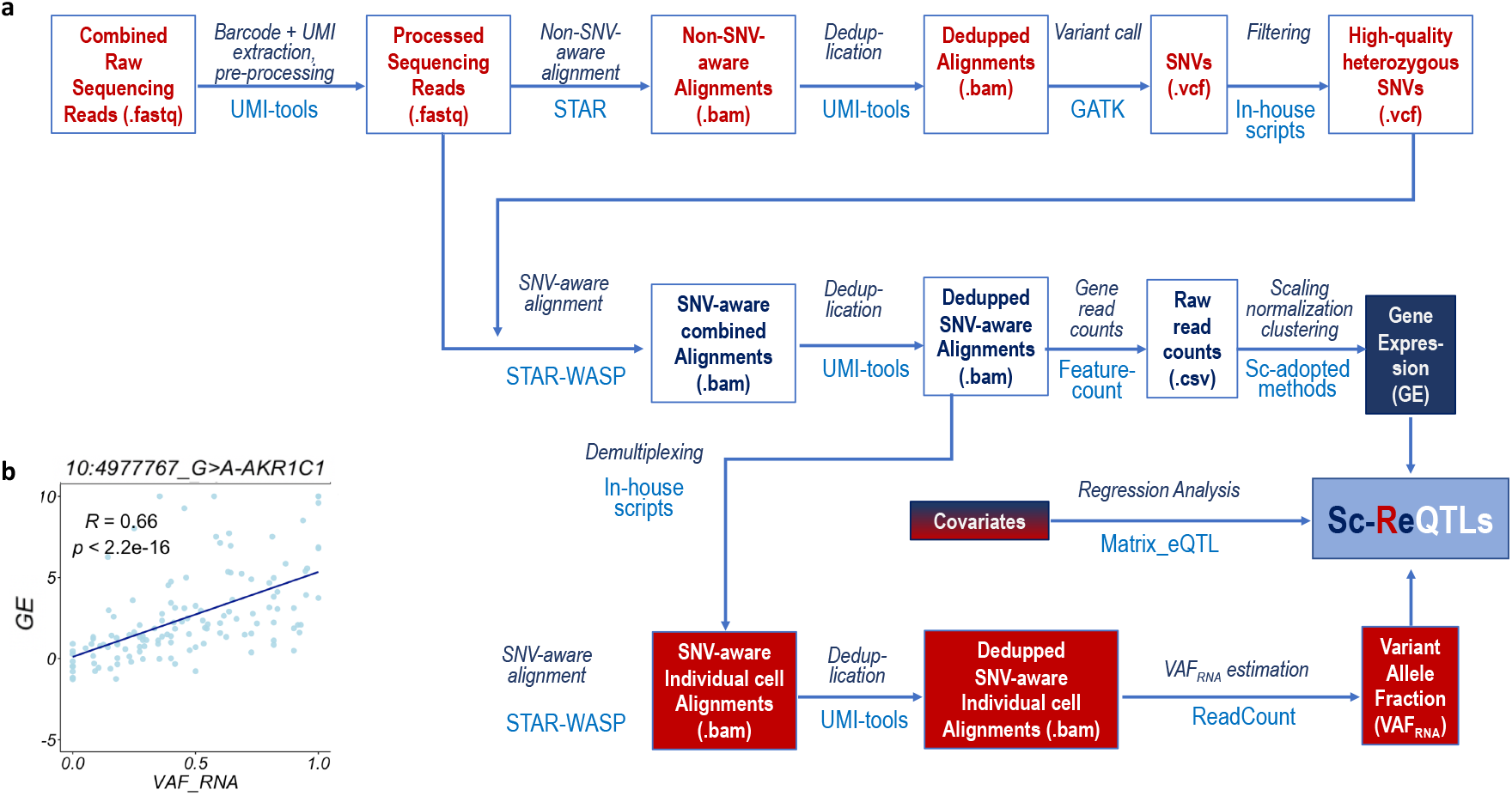
ScReQTL workflow (**a**), with an example of a significant scReQTL correlation between the SNV at 10:4977767_G>A and the gene *AKR1C1* (**b**).

*Processing* includes alignment, deduplication, and variant calling. Because VAF_RNA_ estimations can be sensitive to allele mapping bias, SNV-aware alignment is strongly recommended for VAF_RNA_ – based pipelines. We perform SNV-aware alignment as previously described (26) using 2-pass STAR-WASP (27,28), with intermediate deduplication (UMI-tools, (29)) and variation call (GATK (18)). To outline heterozygous SNV positions for VAF_RNA_ assessment, we apply a series of filtering steps (See Materials and Methods). The filtered SNV sites (per donor) are then used as an input to the second pass, SNV-aware alignment using STAR-WASP (27,28).

*GE estimation* is performed on the SNV-aware alignments, using FeatureCounts to assess the raw gene counts (30), followed by Seurat for normalization and GE variance stabilization (23,31). The generated GE expression values are then used to remove low quality data, batch effects and cell-cycle effects. The before- and after-filtering distributions of genes and RNA-seq reads, and the effects of batch-correction and cell-cycle effects removal are shown on Figure 2. The most variable genes are then identified and used for the scReQTL analyses.

**Figure 2.**
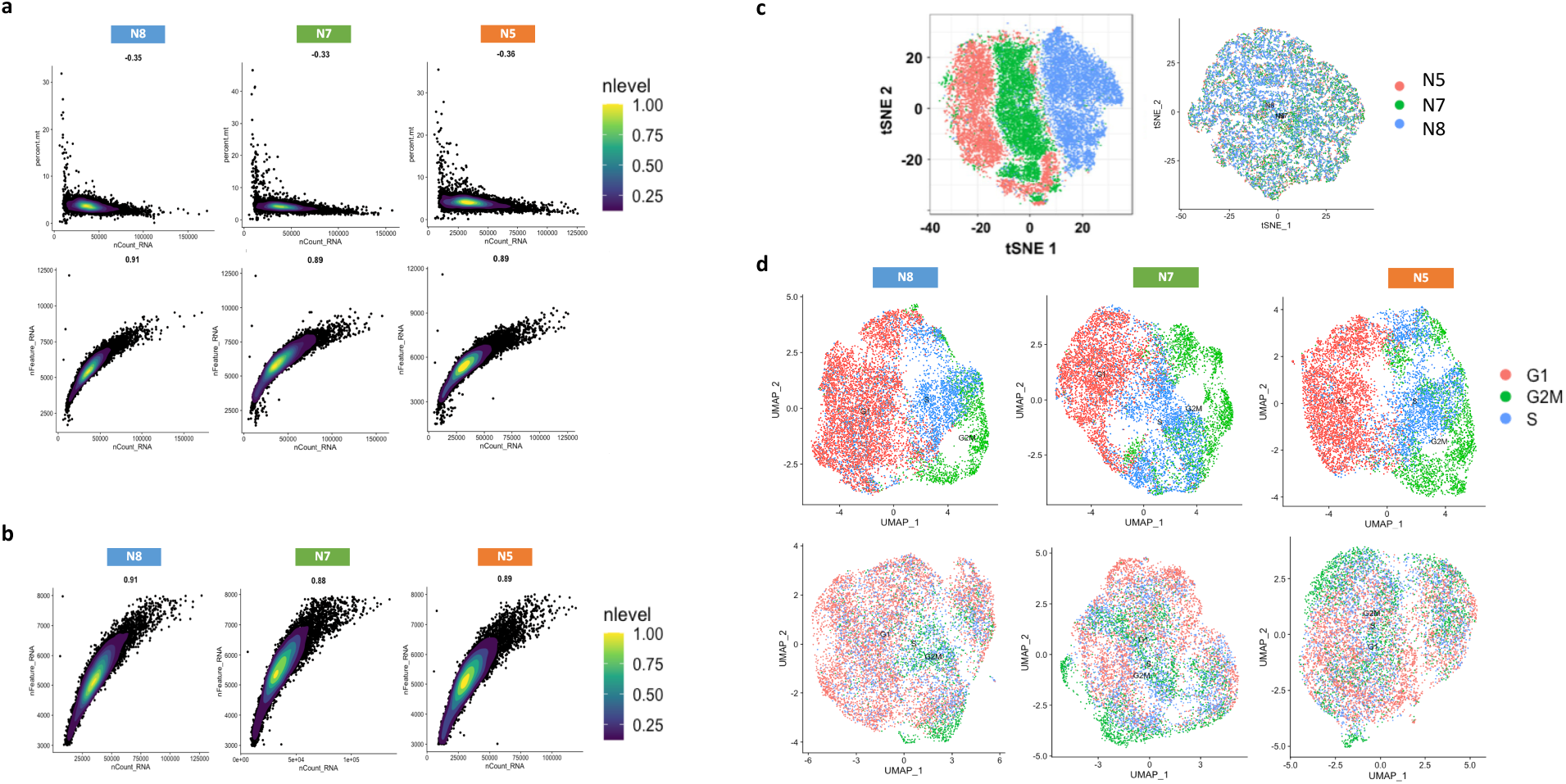
**a**) Density plots showing the proportion of transcripts of mitochondrial origin per cell (top) and the distribution of genes and sequencing reads per cell in the original data (bottom). **b**) Distribution of genes and sequencing reads per cell after filtering of cells with low quality data, defined as less than 3,000 or more than 7,000 genes/cell and/or mitochondrial genes’ expression higher than 6% of the total gene expression. **c**) t-SNE plots before (left) and after (right) correction for batch effects using the Seurat. Strong batch effect are visible before the correction. **d**) Top: cell cycle scores based on expression of G2/M and S phase markers assigned using Seurat. Bottom: Scores after regressing out the cell cycle source of heterogeneity

*Cell type identification* is performed using SingleR (32). The expression profile of each single cell was correlated to expression data from the BluePrint + ENCODE dataset. Across the three study samples, four major cell types were identified: adipose cells, erythrocytes, neutrophils, and naïve-B cells. Adipose cells and erythrocytes were found in all three samples, whereas naïve-B cells were seen in N5 and N7 and neutrophils – in N8 (Figure 3).

**Figure 3.**
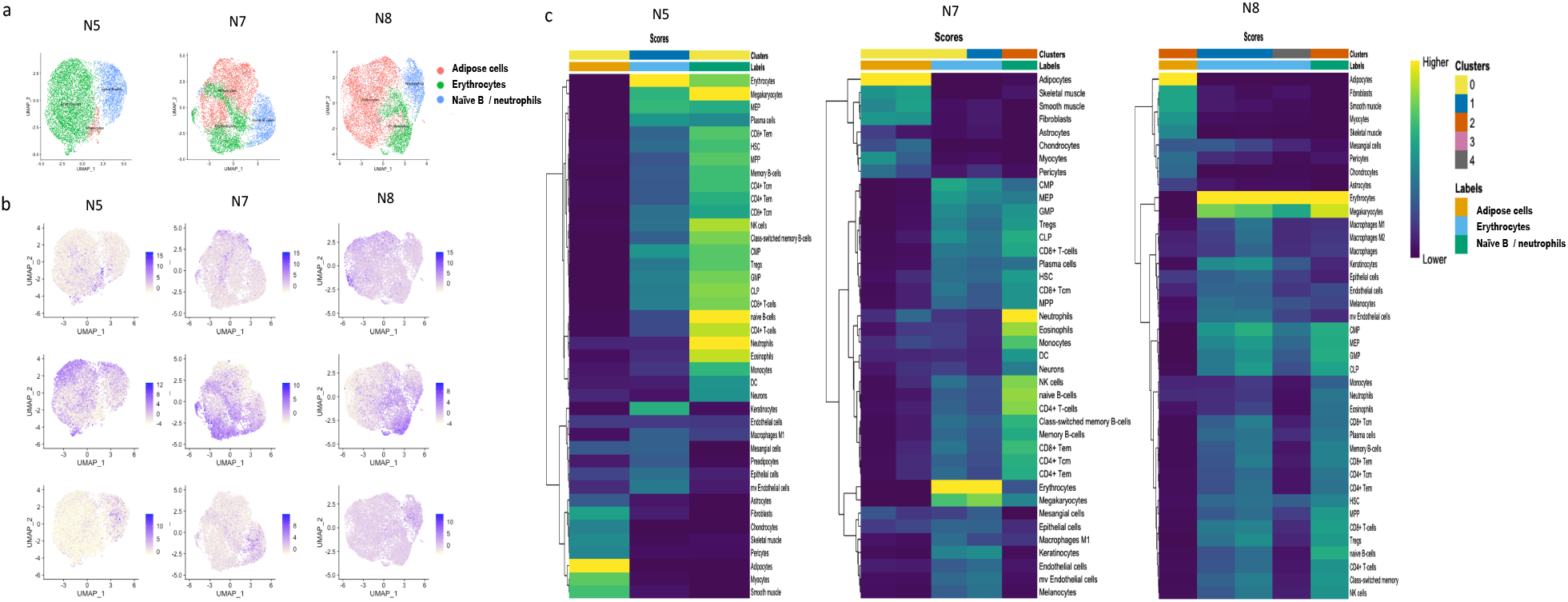
**a**) Cell types identified in each donor using SingleR. Adipose cells and erythrocytes were found in all three donors, whereas naïve-B-cells were seen in N5 and N7 and neutrophils only in N8. **b**) expression of genes associated with cell types: *DCN* (adipose cells, top), *H2AFZ* (erythrocytes, middle), and *H1F0* (neutrophils and naïve B cells). **c**) Heatmap of SingleR scores for top correlated cell types from each of Seurat generated clusters. SingleR uses expression data to regenerate the clusters, and for each cluster, calculates the Spearman coefficient for the genes in the reference dataset. Then, it uses multiple correlation coefficient to collect a single value per cell type per cluster.

*VAF_RNA_* is assessed from the individual cell alignments at sites with heterozygous SNV calls using ReadCounts (22), which estimates the number of sequencing reads harboring the variant and the reference nucleotide (n_var_ and n_ref_, respectively), and calculates VAF_RNA_ (VAF_RNA_ = n_var_ / (n_var_ + n_ref_)) at each heterozygous SNV site of interest (26). To address stochasticity of sampling, estimations of VAF_RNA_ require a threshold of minimal number of unique sequencing reads (minR). Our previous research shows that current scRNA-seq datasets can contain hundreds of SNV sites covered by minimum of 10 sequencing reads (minR ≥ 10) and thousands of SNV sites with minR ≥ 5 (26). In the herein presented analysis, we used VAF_RNA_ estimated at sites with minR ≥ 10; from here on, we refer to these loci as informative. The VAF_RNA_ distribution of the qualifying SNVs is then examined to identify the most variable VAF_RNA_ loci (see Methods). VAF_RNA_ distributions before and after filtering of uninformative (minR<10) and non-variable VAF_RNA_ are shown on Figure 4a and b, respectively.

**Figure 4.**
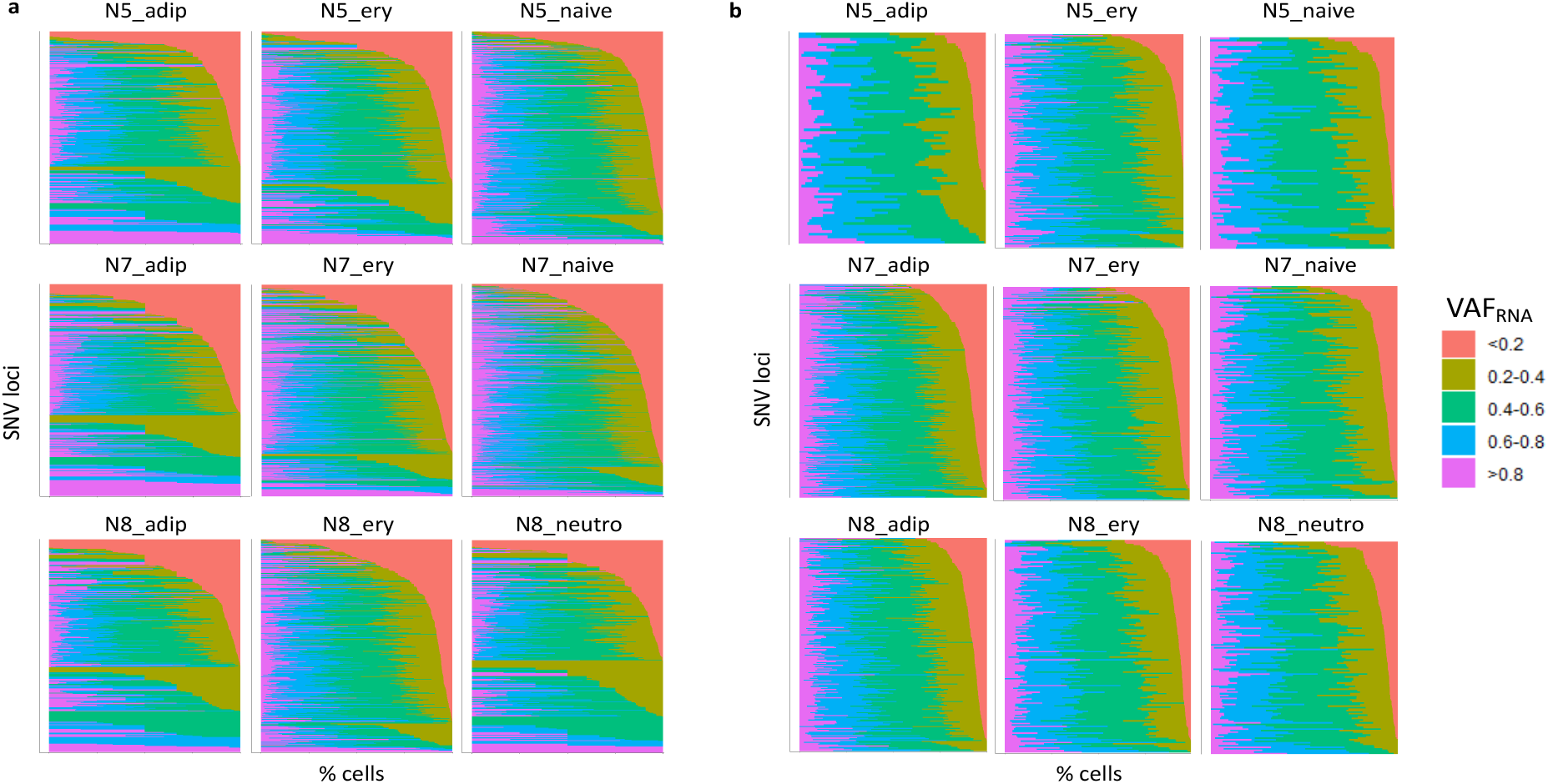
Distribution of scVAF_RNA_ values estimated at SNV sites (displayed on the y-axis) with minR≥10 before (**a**) and after (**b**) filtering of non-variable SNV loci. The SNV sites are sorted by decreasing percentage of cells (x-axis) with scVAF_RNA_ values < 0.2.

*SNV-GE correlations* (scReQTLs) are then computed for each donor, stratified by cell type (see Methods). To qualify for scReQTLs analysis an SNV locus is required to have informative and variable VAF_RNA_ estimations from at least 20 cells per analysis. The variable VAF_RNA_ were correlated to the normalized GE values of the variable genes using linear regression model as implemented in Matrix eQTL (17); quantile-quantile plots (QQ-plots) are presented on Supplementary Figure 1. Cis- and trans correlations were annotated as we have previously described for the bulk ReQTLs (24). Briefly, because scReQTLs are assessed from transcripts, we assign cis-correlation based on the co-location of the SNV locus within the transcribed gene; all the remaining correlations are annotated as trans-scReQTLs).

### Overall scReQTL findings

The number of variable genes and VAF_RNA_ loci retained for scReQTL analysis in the three donors (by cell type) is shown in Table 1. We performed scReQTL analysis separately for each individual and cell type; accordingly, 9 scReQTL analyses were run. Among the samples and cell types, between 79 and 316 SNV loci, and between 2114 and 2442 genes were used as input for scReQTL analysis. Across the 9 groups, a total of 644 distinct SNVs and 2571 distinct genes were tested. This analysis identified 1281 unique scReQTLs at false discovery rate (FDR) of 0.05. All significant scReQTLs are listed in Supplementary Table 1; examples are shown in Figure 5.

**Table 1.** Input parameters for scReQTL analysis, and number of identified scReQTLs per cell type.

| Sample | N cells | N reads | Mean Reads/Cell | Median Genes/Cell | N cells (per cell type) after filtering |  | N input SNVs | N input genes | N scReQTLs FDR = 0.05 |
| --- | --- | --- | --- | --- | --- | --- | --- | --- | --- |
| N5 | 8,906 | 1,071,156,174 | 120,273 | 5,439 | Adipocytes | 296 | 79 | 2,114 | 31 |
|  |  |  |  |  | Erythrocytes | 5,848 | 208 | 2,206 | 161 |
|  |  |  |  |  | Naïve-B cells | 2,033 | 99 | 2,138 | 82 |
| N7 | 8,478 | 1,579,342,505 | 186,287 | 6,049 | Adipocytes | 3,819 | 316 | 2,442 | 336 |
|  |  |  |  |  | Erythrocytes | 2,788 | 238 | 2,395 | 127 |
|  |  |  |  |  | Naïve-B cells | 1,618 | 167 | 2,366 | 102 |
| N8 | 9,256 | 1,285,218,728 | 138,852 | 5,559 | Adipocytes | 5,738 | 230 | 2,345 | 299 |
|  |  |  |  |  | Erythrocytes | 1,924 | 157 | 2,367 | 72 |
|  |  |  |  |  | Neutrophils | 1,433 | 139 | 2,340 | 71 |
| Total/Overall | 26,640 | 3,935,717,407 | 148,471 | 5,682 | Total/Distinct | 25,497 | 1633 / 644 | 20,713 / 2,571 | 1,281 / 1272 |

**Figure 5.**
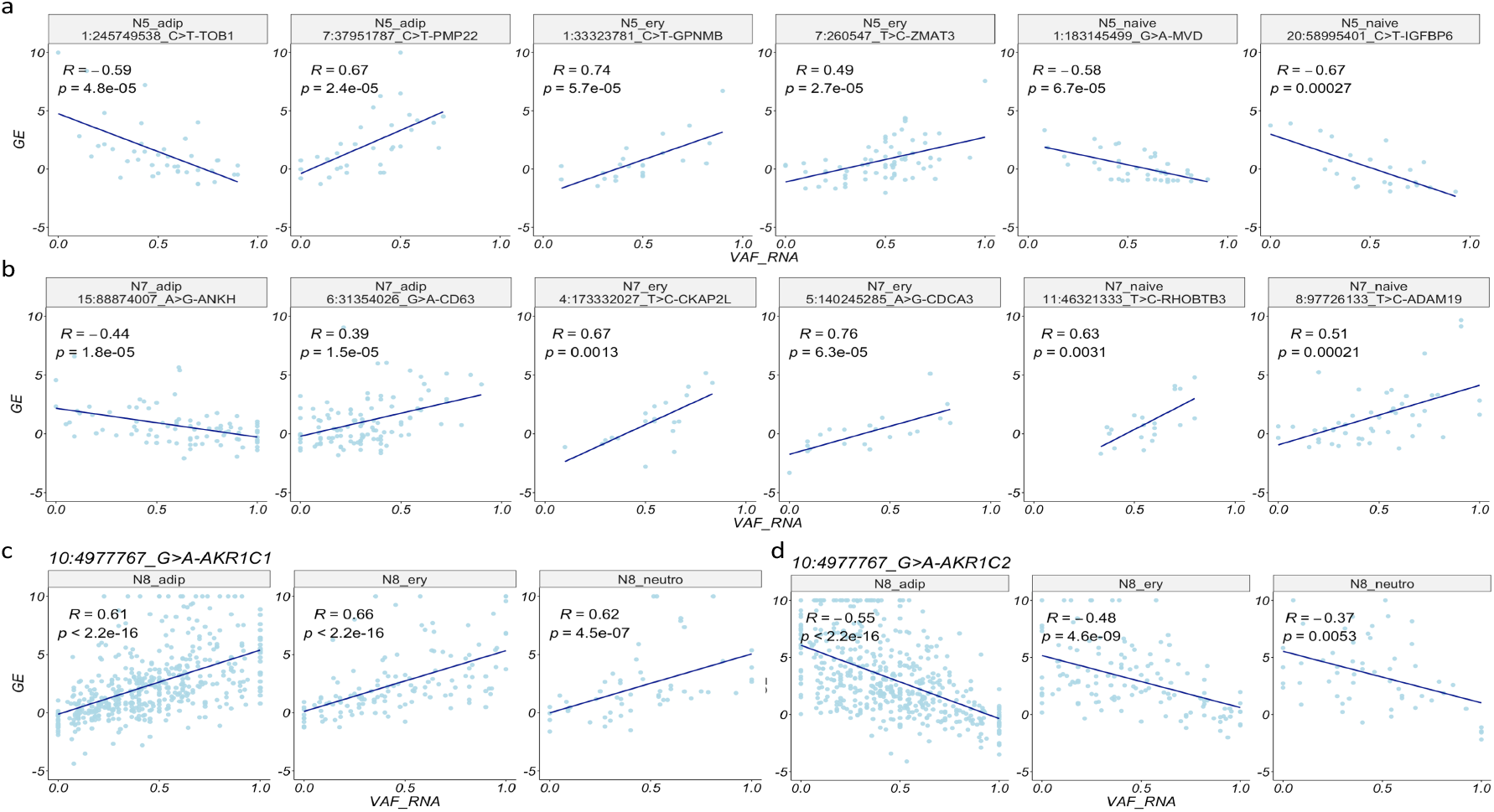
Examples of significant (FDR=0.05) scReQTL correlations in donor N5 (**a**), N7 (**b**) and N8 (**c** and **d**). In N8, consistent across the three cell types cis-scReQTL is shown between the SNV at 10:4977767_G>A and its harboring gene *AKR1C1* (**c**), and between the same SNV and the nearby positioned gene *AKR1C2* (trans-scReQTL, **d**). Note that the displayed P-values are calculated based on the input for the plots generated using the R-package ggplot2 and do not represent the FDR— corrected values from the scReQTL analysis performed with Matrix eQTL.

Among the unique scReQTLs, 7 were identified in more than one cell type or sample (Supplementary Table 2). In all these cases, the correlations were in the same direction, and the effect sizes were similar (See Figure 5c and d). We note that the number of common input SNVs across the 3 samples was as low as 20 (numbers of common input SNVs and genes, as well as the common scReQTLs SNVs and genes are shown in Supplementary Figure 2).

Next, we investigated the relationship between cis- and trans-scReQTLs. Of the significant scReQTLs, only 6 represented cis correlations (See examples in Figure 5c). This observation differs from eQTL analyses, which typically identify a high number of significant cis-correlations. Here we note that the ReQTL annotation of cis- and trans-differs from the distance-based annotation used for eQTLs, which considers cis-regulatory SNVs in nearby genes and transcriptionally silent genomic regions. We then assessed if some scReQTLs are mediated by cis-effects that do not reach significance at an FDR of 0.05. To do this, we computed the correlation of all SNVs represented in significant trans scReQTLs with their harboring gene. For 26% of the scReQTL SNVs, we detected correlations with their harboring genes with 0.05 < FDR < 0.1 (Supplementary Figure 3). This analysis suggests that a proportion of the SNVs may at least partially exert their trans-effects via weak to moderate regulation of the expression of their harboring gene.

### scReQTL in known genetic networks

To assess to what extend scReQTL findings agree with known SNV-gene, and gene-gene interactions, we intersected the significant scReQTLs with: (a) eQTLs reported in the GTEx database (8), (b) ReQTLs as estimated from bulk adipose sequencing data (24), (c) known gene-gene interaction from the STRING database (33), and (d) significant GWAS loci (34).

#### scReQTLs and eQTLs from GTEx

To estimate the overlap between scReQTL and known eQTLs, we used the data from 49 different tissues and cell types from the GTEx database (https://www.gtexportal.org). First, we identified the SNVs and genes used as an input for scReQTLs, and participating in known eQTLs: a total of 111 input SNVs and 2024 input genes participated in at least one eQTL reported in GTEx. Across the 49 tissues, scReQTL identified 32 correlations (Supplementary Table 3), comprised of 6 unique SNV-gene pairs (5 SNVs and 6 genes). These pairs included all 4 significant cis-scReQTLs, and two trans-scReQTLs: chr10_4977767_G>A and *AKR1C2* (see Figure 5d), and chr1:115337511_G_A and *NGF*. For each of the 6 SNV-gene pairs, we compared the scReQTLs and the eQTLs in the different GTEx tissue types. For 3 of the 6 scReQTLs, the corresponding GTEx eQTLs were consistent in terms of directionality and effect size (Figure 6 and Supplementary Figures 4 and 5).

**Figure 6.**
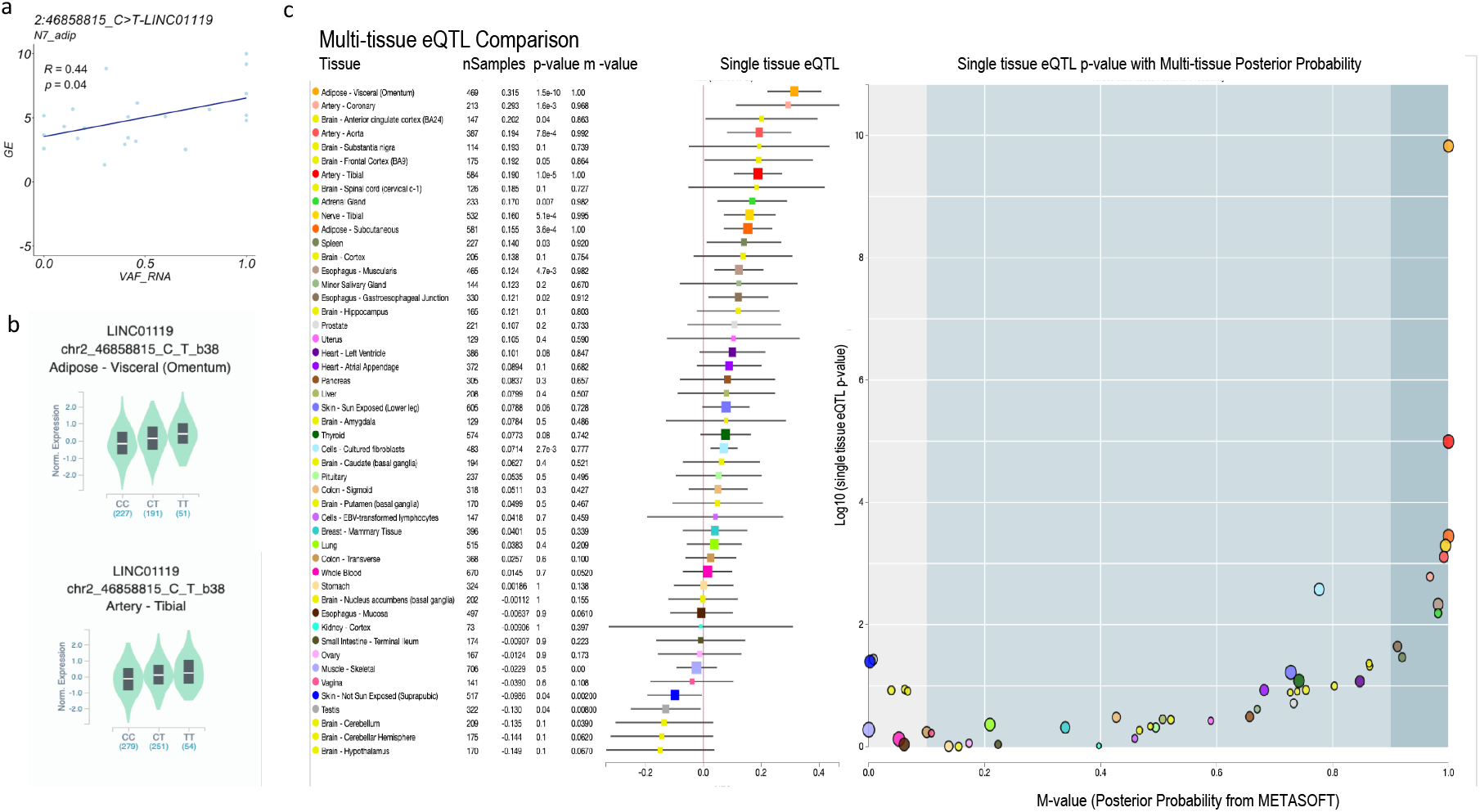
scReQTL and eQTLs between the SNV at 2:46858815_C>T and its harboring gene *LINC01119* (cis-scReQTL). **a**) scReQTL between the SNV at 2:46858815_C>T and *LINC01119*. **b**) eQTLs between the SNV 2:46858815_C>T and *LINC01119* reported in the GTEX in 2 tissues: Adipose Visceral and Artery – Tibial. The graphs are generated at the GTEx portal (https://www.gtexportal.org/). The eQTLs and scReQTL agreed in terms of directionality and effect sizes. **c**) Multi-tissue comparisons of the eQTL at 2:46858815_C>T and *LINC01119* generated at the GTEx portal (https://www.gtexportal.org/).

The other 3 scReQTL were found as both positive and negative eQTLs depending on the tissue type in GTEx. The positive cis-scReQTL, chr6:31354105_G>A_HLA-B, was a significant cis-eQTL in 4 GTEx tissues: positive in three, but negative in the testis (Supplementary Figure 6). The last 2 scReQTLs comprised correlations of the SNV at chr10:4977767_G>A with *AKR1C1* (positive) and *AKR1C2* (negative); these scReQTLs were consistent across cell types (see Figure 5c and d). In GTEx, the corresponding eQTLs were found in multiple tissues, and in both positive and negative correlations, highlighting tissue-specific effects (Supplementary Figures 7 and 8).

Overall, our analysis on the agreement between significant scReQTLs and eQTLs identified a narrow overlap, within which most observations were consistent, and the remaining were not contradictory. We note that this analysis was limited by the relatively small number of input scReQTL SNVs present in GTEx. Furthermore, while the cis-scReQTLs agreed with the cis-eQTLs, the majority of the significant scReQTLs were in trans, which are known to be highly tissue-specific (8). None of the 4 cell types assessed in our study - adipose cells, erythrocytes, neutrophils, and naïve-B cells obtained from adipose-derived mesenchymal stem cells - were a direct match to any of the 49 tissues and cell types from the GTEx database. Finally, we expect that the strongest contributor to the low level of concordance between scReQTL and eQTLs is the limited detection power of scReQTL due to the sparsity of the scRNA-seq data, which is reflected in the low number of cells passing the minR requirement for each SNV locus and included in the regression analysis. Indeed, while the initial cell counts per scReQTL analysis (except for N5 adipose cells) were over 1000, the majority of the SNV loci had between 20 (the required minimum) and 100 cells with minR>10 per cell type (Supplementary Figure 9a). In comparison, the GTEx eQTLs are computed from a minimum of 100, and in most of the tissues, from over 250 individuals (Supplementary Figure 9b).

#### scReQTLs and ReQTLs from bulk adipose tissue

Next, we intersected the scReQTL findings with ReQTLs from bulk RNA-sequencing data. To do this, we performed ReQTL on RNA-seq data from two adipose tissues downloaded from GTEx – adipose subcutaneous (275 samples) and adipose visceral (215 samples) - following the published protocol (24). Using the SNVs and the genes used as input for the scReQTL, with an FDR = 0.05, ReQTL did not identify significant correlations, whereas with an FDR = 0.1, ReQTL identified 84 (6.6%) and 48 (3.8%) of the significant scReQTLs, in adipose subcutaneous and visceral tissue, respectively. The majority of the these ReQTLs had small effect sizes and agreed in the direction with the corresponding scReQTL in 71% of the cases (Examples shown on Figure 7a). Of note, the above discussed chr10:4977767_G>A and *AKR1C1*/*AKR1C2* did not show any correlation when examined from bulk RNA-seq data (Figure 7b).

**Figure 7.**
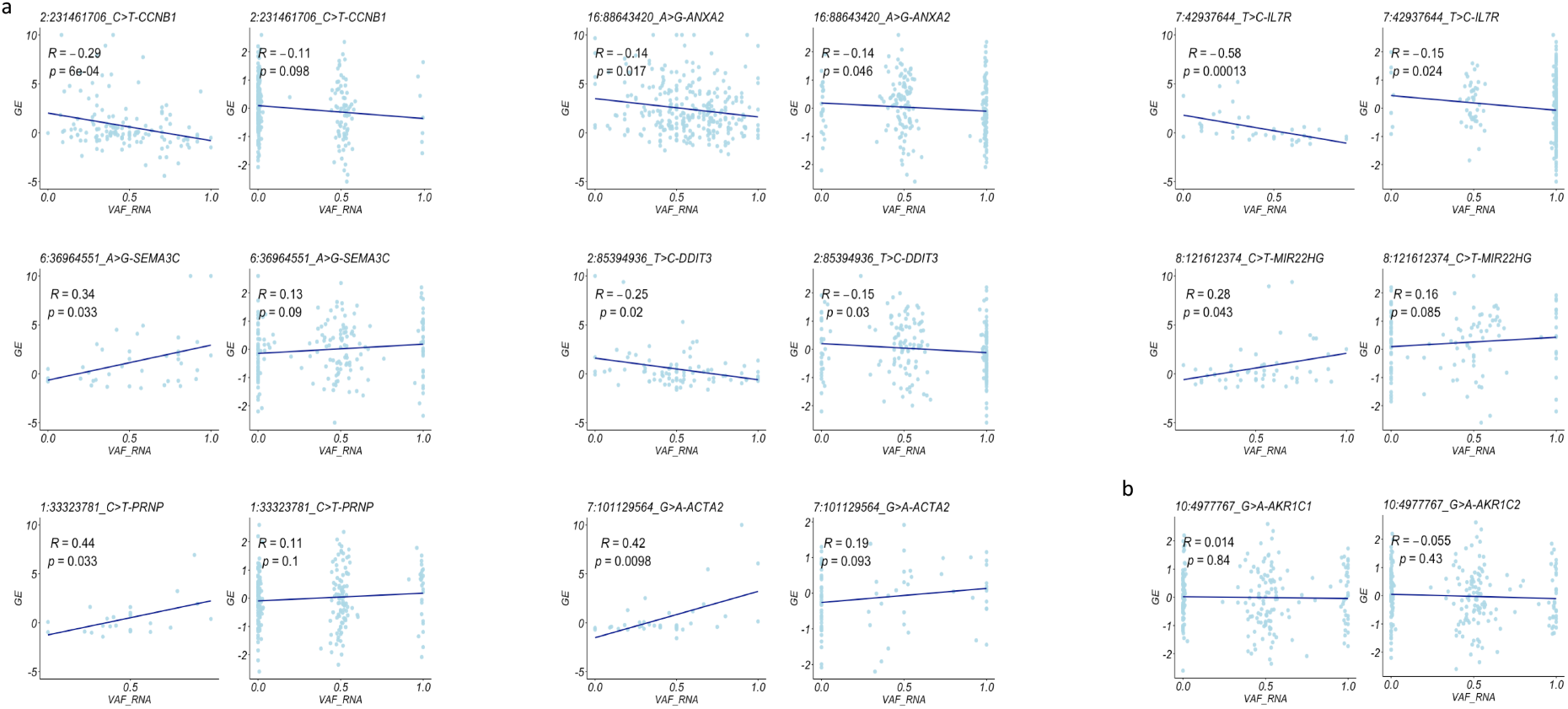
scReQTLs and ReQTLs from bulk adipose tissue. **a**) Examples of comparisons of scReQTLs (left) and ReQTLs from bulk adipose tissue (right) at FDR = 0.1. The ReQTLs had generally weaker size effects and agreed in directionality in 71% of the correlations. Note that the displayed P-values are calculated based on the input for the plots generated using the R-package ggplot2 and do not represent the FDR—corrected values from the scReQTL analysis performed with Matrix eQTL. b) ReQTL analysis between the SNV at 10:4977767 and *AKR1C1* (left), and *AKR1C2* (right), which were found as significant scReQTLs, did not show significant correlation in bulk RNA-seq data.

The lack of strong overlap between scReQTL and ReQTL (as well as eQTL) suggests different regulatory relationships captured by scReQTLs. While ReQTLs and eQTLs show a high overlap between each other, and are both based on abundance of variant alleles across multiple individuals with different genotypes, scReQTL operates in a setting of identical genotypes, and reflects cell-specific networks that are likely to capture transient, allele-mediated genetic interactions.

#### scReQTLs and known gene-gene interactions

Because the vast majority of the significant scReQTLs reflected correlations between two different genes (VAF_RNA_ of one of the genes and expression level of the other), we assessed if these gene pairs were enriched in known gene-gene interactions. We downloaded the known gene-gene (human) interactions from the STRING database (33) and intersected these with the scReQTLs. From the 1234 unique gene-gene scReQTLs pairs, 203 (16.4%) were previously annotated in STRING (Supplementary Table 4, p < 10e-4, permutation test using 10000 permutations, Figure 8a). Examples include *IFIT1* and *IFITM2*, *AURKA* and *PLK, and CKS2* and *CDC20* (Figure 8b-c). The strong enrichment of scReQTLs with known genetic networks suggests that scReQTLs may be used to identify allele contributions to gene-gene interactions.

**Figure 8.**
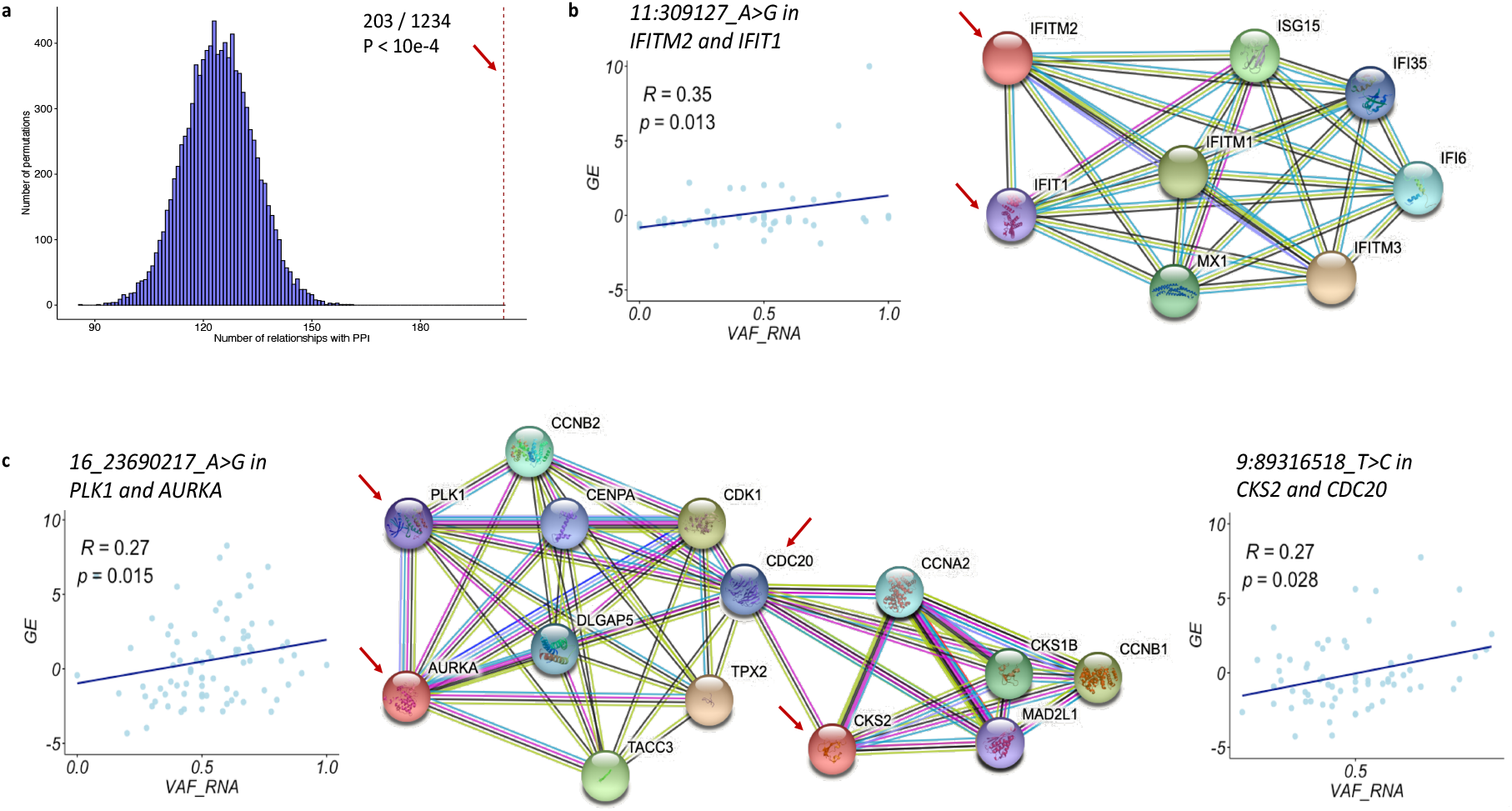
**a**) Permutation test for assessment of enrichment of trans scReQTLs in known gene-gene interactions obtained from the STRING database; 10000 permutations were used. The p-value (p<10e-4) was defined as the fraction of permutations in which the number of gene-gene pairs found in the known interaction database was at least as great as the number found in the observed data. This analysis showed significant enrichment of trans-scReQTLs with known gene-gene interactions. **b** and **c**) Examples of trans-scReQTLs and known gene-gene interactions: *IFITM2* (11:309127_A>G) in and *IFIT1* (b) and *PLK1* (16_23690217_A>G) and *AURKA*, and CKS2 (9:89316518_T>C) and CDC20 (**d**). The interaction graphs are generated using the STRING database visualization tools. Note that all the scReQTL highlighted gene-gene interactions are supported by a minimum of three lines of evidence that include either experimental validation (purple line) or curated databases (light-blue line), or both.

#### scReQTLs and GWAS

Furthermore, we intersected the SNVs participating in scReQTLs with SNVs significantly associated with phenotypes by GWAS (35). This analysis showed that 18 (out of the 408 unique scReQTL SNVs, 4.4%) were present in GWAS; these 18 SNVs participated in 84 scReQTL correlations (Supplementary Table 5). This percentage is similar to the overlap between GWAS and GTEx eQTLs (3.7 and 3.6% in adipose visceral and adipose subcutaneous tissue respectively), and significantly higher than the overlap with common SNVs from DbSNPv.154, (0.34%, p < 10e-6). This analysis shows that scReQTL SNVs are enriched in genetic variants associated with phenotype via large population-based and case-control studies.

### Functional scReQTLs SNVs annotations

We assessed the SNVs participating in scReQTL in regards to position in the harboring gene and predicted functional effects. As expected from scRNA-seq data generated using a 3’-based protocol, the majority of the SNVs resided in the 3’UTR of their harboring gene (70.2%, Supplementary Figure 10); the 3’UTR SNVs participated in 69.6% of the scReQTLs. 3’-UTR variants are known to strongly affect both GE levels and splicing (36–39); hence, scReQTLs can be applied to study this aspect of genetic regulation. The second category was exonic SNVs, comprising 16.2% of the unique SNVs and participating in 14.9% of the scReQTLs. Exonic SNVs included missense, nonsense, and near-splice variants, many of which can potentially affect the protein structure and function. Of note, scReQTL captured a substantial number of intronic SNVs – 13%, participating in 11.2% of the scReQTLs. Intronic sequences are reported in 15%–25% of the RNA-sequencing reads from both bulk and single-cell RNA-seq (4,38,39). Intron quantitation can be used to estimate the relative abundance of precursor and mature mRNA, thereby assessing the RNA velocity and dynamic cellular processes (4). In the allele-specific setting provided by the scReQTLs, correlation of intronic SNVs with GE can identify SNVs regulating the RNA processing and maturation.

Next, we assessed if the scReQTLs SNVs are enriched in specific clinical phenotypes obtained from the ClinVar database (40). Fifteen SNVs (3.7% of the total 408 distinct scReQTL SNVs) were associated with known clinical phenotypes, including circulating phospholipid trans fatty acids, cortisol levels, circadian rhythm, risk for cardiovascular disease, blood pressure, schizophrenia, neuroticism, osteoporosis, anthropometric traits, and asthma (See Supplementary Table 1). This percentage is similar to the overlap between ClinVar and GTEx eQTLs (3.3 and 3.1% of the eQTLs in adipose visceral and adipose subcutaneous tissue respectively), and significantly higher than the overlap with common SNVs from DbSNPv.154, (0.61%, p < 10e-6). Finally, we assessed the predicted functional and/or pathogenic scores of the scReQTL SNVs using 17 models including SIFT, Polyphen2, LRT, MutationTaster, MutationAssessor, FATHMM, PROVEAN, VEST3, CADD, DANN, fathmm-MKL, MetaSVM, MetaLR, integratedFit, GERP++, phyloP, and phastCons, as implemented in ANNOVAR (41); this data is summarized in Supplementary Table 6).

### scReQTL application

Application of scReQTLs requires consideration of several factors. First, because scReQTLs are confined to expressed SNV loci, they cannot capture variants in transcriptionally silent genomic regions. In addition, SNV loci with expression levels below the required minimum number of RNA-seq reads (minR) are not included in the scReQTL analyses. Furthermore, because of the platform used in this study - 10× Genomics Chromium v3 chemistry – the analyzed SNVs are restricted to those located within the length of the sequencing read (here, 150nt) from the 3’ end of the transcript. For many genes, these reads cover only a proportion of the SNVs residing in a transcript. For the above reasons, scReQTLs accessible SNVs represent a relatively small subset of the expressed SNVs and are not designed to cover the full set of SNVs in the transcriptome.

Second, it is important to note that even when a genetically regulated gene is captured by scReQTL analysis, the scReQTLs may not include the actual causative SNV, but its co-allelic SNVs. This is the case for SNVs positioned outside the transcribed regions or outside the coverage of the sequencing library.

Third, scReQTLs are based on VAF_RNA_ estimation, which can be affected by technical parameters, including allele mapping bias (42) which can lead to overestimation of the reference allele count (43). Therefore, we perform the scReQTL using SNV-aware alignments. Specifically, we apply STAR-alignment with WASP, which removes ambiguously mapped reads after checking for consistency with the reads containing the alternative nucleotide (27,28).

Another important parameter for VAF_RNA_ estimation is the selection of cutoff for minimal number of reads, minR. When selecting minR for an analysis, a major factor is the balance between the confidence of VAF_RNA_ estimation (high minR) and the inclusivity of SNVs (lower minR values include more loci for scReQTL). In the present study, we have included SNV loci with minR ≥ 10. Our previous research shows that for current 10× Genomics scRNA-seq datasets, minR ≥ 5 provides a reasonable balance between VAF_RNA_ confidence and SNV inclusivity (26). At lower cutoffs (i.e. minR = 3) stochasticity of sampling can affect the VAF_RNA_ estimation (26). In addition, low cutoffs are expected to include SNVs in genes expressed at low levels, where additional technical noise can affect the accuracy of the estimations.

Furthermore, VAF_RNA_ can be affected by inaccuracies in the variant calling, including incorrect calling of the presence or absence of an SNV, and erroneous assignment of a heterozygous state. The presented pipeline uses scRNA-seq data only, where we call SNVs from pooled scRNA-seq data, and select for scReQTL analysis highly confident heterozygous sites based on mapping and Phred quality, genomic position (genic, non-repetitive regions), and previously validated rsID. To confidently assign heterozygosity, we select bi-allelic SNVs with a minimum of 50 unique reads supporting each allele from the pooled scRNA-seq. By default, this selection excludes heterozygous SNVs with strong non-random monoallelic expression. Therefore, while the above approach is suitable for datasets where matched DNA is not available, we recommend assignment of heterozygosity based on genotypes when available. Importantly, scReQTLs do not necessarily require prior variant calls and can be run on custom pre-defined lists of genomic positions such as dbSNP or a database of RNA-editing sites.

Finally, VAF_RNA_ varies between different cell types, often due to cell-specific regulatory mechanisms (44). Due to the dynamic nature of RNA transcription, it is expected that VAF_RNA_ (similarly to GE) will vary depending on conditions, disease states and stochastic factors. Therefore, interpretation of scReQTL findings requires consideration of the dynamics of the variables underlying the correlation.

## Discussion

Single-cell RNA-seq eQTL analyses define an emerging research niche that brings major benefits for the understanding of functional genetic variation including the identification of cell-type and condition-specific correlations (2,13–16,45). In this paper, we present a new eQTL-based analysis in a scRNA-seq setting - scReQTL – which uses the VAF_RNA_ at expressed heterozygous SNVs in place of the genotypes, to correlate allele prevalence to gene-expression levels. By using VAF_RNA_ across multiple cells of the same sample, scReQTLs introduce several new analytical aspects.

First, and perhaps most importantly, as scReQTL can be implemented on multiple single cells from the same sample, it can be applied to assess the effects of SNVs in a single sample or individual. This is particularly applicable for rare SNVs which are challenging to study via population-based approaches. Second, scReQTLs increase the dynamicity of the SNV-gene correlations, as VAF_RNA_, similarly to GE, is both dynamic and cell-type-specific (44). In particular, in each cell type, scReQTL correlates the most variable VAF_RNA_ to the most variable genes. Third, as compared to the discrete genotype values (0,1,2), VAF_RNA_ can obtain continuous values spread along the entire VAF_RNA_ range ([0,1]), allowing for more precise computation of the proportion of each allele represented in the RNA in a given cell. Fourth, scReQTL operates in the context of (largely) identical genotypes, which narrows the observed effects to RNA-mediated interactions. Finally, scReQTL does not necessarily require matched DNA (although we recommend it for genotyping of heterozygous SNVs, if available), and therefore can be applied on scRNA-seq data alone. Related to that, scReQTL analyses can be performed using pre-defined SNV lists, such as RNA-editing sites and sets of dbSNP SNVs of interest.

At the same time, compared to single cell and bulk eQTLs, scReQTL analyses have notable limitations. First, the scReQTL accessible SNVs are restricted by depth of coverage per cell (minR) and, in the case of 3’-based scRNA-seq protocols, by the length of the sequencing read. Therefore, scReQTLs can analyze only a proportion of the transcribed SNVs. This limitation is expected to be gradually reduced with the progress of the sequencing technologies. Additional attenuation of this constraint is possible through reducing the value of minR used in the analysis. Indeed, while in this study we apply minR ≥ 10, which retained between 308 and 721 input SNVs per sample, in our prior research we show that at minR ≥ 5 the number of SNVs is higher by an order of magnitude (26). Second, scReQTL appears to have relatively low power to detect cis-acting (on the same gene) SNVs (See Supplementary Figure 3). Specifically, the vast majority of the correlations identified in this study are trans-scReQTLs. Several factors may account for this observation. As mentioned earlier, the definition of “cis”-scReQTLs is based on residing of the SNV within the same gene; hence SNVs that would be classified as “cis” using the eQTL distance-based definition are “trans” for the scReQTLs, increasing the proportion of trans-correlations in the same SNV-gene dataset. Additional possible explanation is that in the explored setting of minR≥10, cis-acting SNVs are located in genes with high expression, which likely contain a high proportion of stably expressed genes, including with house-keeping functions. Confining the analyses to SNVs in genes with high expression level is an additional limitation of the scReQTLs. Nevertheless, due to the dynamic nature of the scReQTL estimations, scReQTLs can capture SNVs in genes with transiently high expression in a particular cell type or in a specific stage of the cell development. Notably, the identified trans-scReQTLs are significantly enriched in known gene-gene correlations (See Figure 7), therefore we interpret them as indictive of an allelic contribution to these gene-gene interactions. The above limitations, together with the relatively low number of cells with minR ≥10 for many of the participating SNVs, at least partially account for the narrow overlap between scReQTLs and eQTLs/ReQTLs. At the same time, scReQTLs are able to capture correlations that are masked in the bulk eQTL and ReQTL analyses (See Figure 8).

Our scReQTL analysis includes approximately 4 billion RNA-seq reads from 26,640 human adipose-derived mesenchymal stem cells, obtained from three healthy donors. We chose the 10×Genomics platform due to its growing popularity, high throughput, and the support for unique molecular identifiers (UMI) for the removal of PCR-related sequencing bias. Using stringent cutoff for SNV coverage (minR≥10) we identified 1272 distinct scReQTLs. These scReQTLs include a considerable number of correlations which involve SNVs previously highlighted by GWAS and are significantly enriched in known gene-gene interactions. These results demonstrate that scReQTLs can be used to identify novel genetic interactions, including those which are specific to a given cell-type.

## Conclusion

We present a new approach – scReQTL – that correlates SNVs to gene expression from scRNA-seq data. The scReQTL analyses presented in this research generated results containing both previously known and novel genetic interactions. scReQTL is applicable to the rapidly growing source of scRNA-seq data, and is capable of identify SNVs contributing to cell type-specific intracellular genetic interactions.

## Materials and Methods

### Data

We used publicly available scRNA-seq data (25) from 26,640 human cells from three healthy donors: N5, N7 and N8. The scRNA-seq data was generated on 10× Genomics Chromium v2 platform; the library preparation and sequencing are described in detail elsewhere (25). Briefly, cells were partitioned using 10× Genomics Single Cell 3’ Chips, and barcodes to index cells (16bp) and transcripts (10bp UMI) were incorporated. The constructed libraries were sequenced on an Illumina NovaSeq 6000 System in 2×150bp paired-end mode.

### SNV-aware alignment

The cell barcodes and UMIs were extracted using UMI-tools from the pooled (per donor) raw sequencing reads (29). The pooled sequencing reads were aligned to the latest version of the human genome reference (GRCh38, Dec 2013) using STAR v.2.7.3.c in 2-pass mode with transcript annotations from the assembly GRCh38.79 (27). The alignments were deduplicated retaining the reads with the highest alignment scores (29). SNVs were called in the pooled deduplicated alignments using GATK v.4.1.4.1 (18). To identify heterozygous SNV positions qualified for VAF_RNA_ analysis, we applied a series of filtering steps. Specifically, heterozygous SNVs were selected based on the presence of minimum of 50 high-quality reads supporting both (reference and alternative) nucleotides in the pooled alignments. SNV loci were annotated using SeattleSeq v.13.00 (dbSNP build 153), and loci positioned in repetitive or intergenic regions were removed. The SNV lists were further filtered based on the following requirements: QUAL (Phred-scaled probability) > 100, MQ (mapping quality) > 60, QD (quality by depth) > 2, and FS (Fisher’s exact test estimated strand bias) = 0.000. The filtered SNV lists (per donor) were then used as an input for a second, SNV-aware alignment using STAR-WASP (28).

### Gene Expression estimation

To estimate gene expression, we first apply FeatureCount on the individual alignments to assess the row gene counts per cell (30). We then normalize and scale the expression data using the *sctransform* package as implemented in Seurat v.3.0 (23,31), which stabilizes the GE variance using regularized negative binomial regression. The normalized GE values are then used to remove cells with low quality data, defined as less than 3,000 or more than 7,000 detected genes and/or mitochondrial genes’ expression higher than 6% of the total gene expression. The GE values were used to correct for batch- and cell-cycle effects (See Figure 2). Thereby selected most variable genes were then used to classify cell types (See below). In addition, after examining the GE distribution across the cells (per cell type), genes which expression in 80% or more of the cells was within 20% or less from the top or bottom of the GE range, were filtered out; the retained most variable genes were then used for scReQTL analyses (See Table 1).

### Cell type identification

To define individual cell types, we used SingleR version 1.0.5 (32). The expression profile of each single cell was correlated to expression data from the BluePrint + ENCODE dataset, containing 259 bulk RNAseq samples representing 24 main cell types and 43 subtypes. SingleR first calculates a Spearman coefficient for the correlation of the expression of the most variable genes of each single-cell gene with each of the samples in the reference data set. Then, it uses multiple correlation coefficient to collect a single value per cell type per cluster. This correlation analysis is rerun iteratively using only the top cell types from the previous step and the variable genes among them until only one cell type is retained. Applying SingleR, we identified four major cell types were identified across the three donors: adipose cells and erythrocytes were found in all three samples, naïve-B-cells found in N5 and N7, and neutrophils, in N8 (See Figure 3 and Table 1).

### VAF_RNA_ estimation

VAF_RNA_ is assessed from the individual alignments as we have previously described (26), using the high quality heterozygous SNV sites as inputs for ReadCounts (22). At each position of interest, ReadCounts estimates the number of sequencing reads harboring the variant and the reference nucleotide (n_var_ and n_ref_, respectively), calculates VAF_RNA_ (VAF_RNA_ = n_var_ / (n_var_ + n_ref_), and filters out positions not covered by the user-defined minimum number of reads (minR); minR is constant across the genome (22). For the herein presented analysis, we used minR>10. To qualify for scReQTL, a variant is required to have variable VAF_RNA_ from a minimum of 20 cells from the same cell type (per donor). The VAF_RNA_ distribution is then examined and loci with non-variable VAF_RNA_ are filtered out. Loci were considered non-variable if: (1) over 75% of the VAF_RNA_ values are in the range of 0.5 ± 0.1 (corresponding to stable biallelic expression), and (2) over 75% of the VAF_RNA_ values are in the ranges 0-0.25 or 0.75-1 (corresponding to predominantly monoallelic or skewed allelic expression).

### ScReQTL computations

*SNV-GE correlations* (scReQTLs) were computed for each donor, across the cells of each type separately. To qualify for scReQTLs analysis, an SNV locus is required to have informative and variable VAF_RNA_ estimations (minR≥10) from at least 20 cells per analysis. The variable VAF_RNA_ were correlated to the normalized GE values of the most variable genes using a linear regression model as implemented in Matrix eQTL (17). The top 15 principal components of the GE were used as covariates (Supplementary Figure 11). Cis and trans correlations were annotated as previously described for the bulk ReQTLs (24). Briefly, because scReQTLs are assessed from transcripts, we assign cis-correlation based on the co-location of the SNV locus within the transcribed gene, using the gene coordinates (46). All the scReQTLs including SNVs residing in genes different from the expression-correlated genes are annotated as trans-scReQTLs.

### Statistical Analyses

Throughout the analysis we used the default statistical tests (with built-in multiple testing corrections) implemented in the used software packages (Seurat, SingleR, Matrix eQTL), where p-value of 0.05 was considered significant, unless otherwise stated. For estimation of differences in overlap between scReQTL SNVs, GWAS and ClinVar, chi-square test was used. For assessment of enrichment of scReQTLs in known gene-gene interactions, a permutation test with 10000 permutations was applied. For each permutation, a random set of gene-gene pairs of the same size as the observed data was selected. The p-value was defined as the fraction of permutations in which the number of gene-gene pairs found in the known interaction database was at least as great as the number found in the observed data.

## Author Contributions

AH: Conceptualization, analysis, writing and supervision; HL, NMP, LS, PB, NA, HI, JS, PS, KTA data management and processing, analysis, visualization, and writing—review and editing.

