## Supplementary figures and images for "scReQTL: an approach to correlate SNVs to gene expression from individual scRNA-seq datasets"

### Supplementary Figure 1

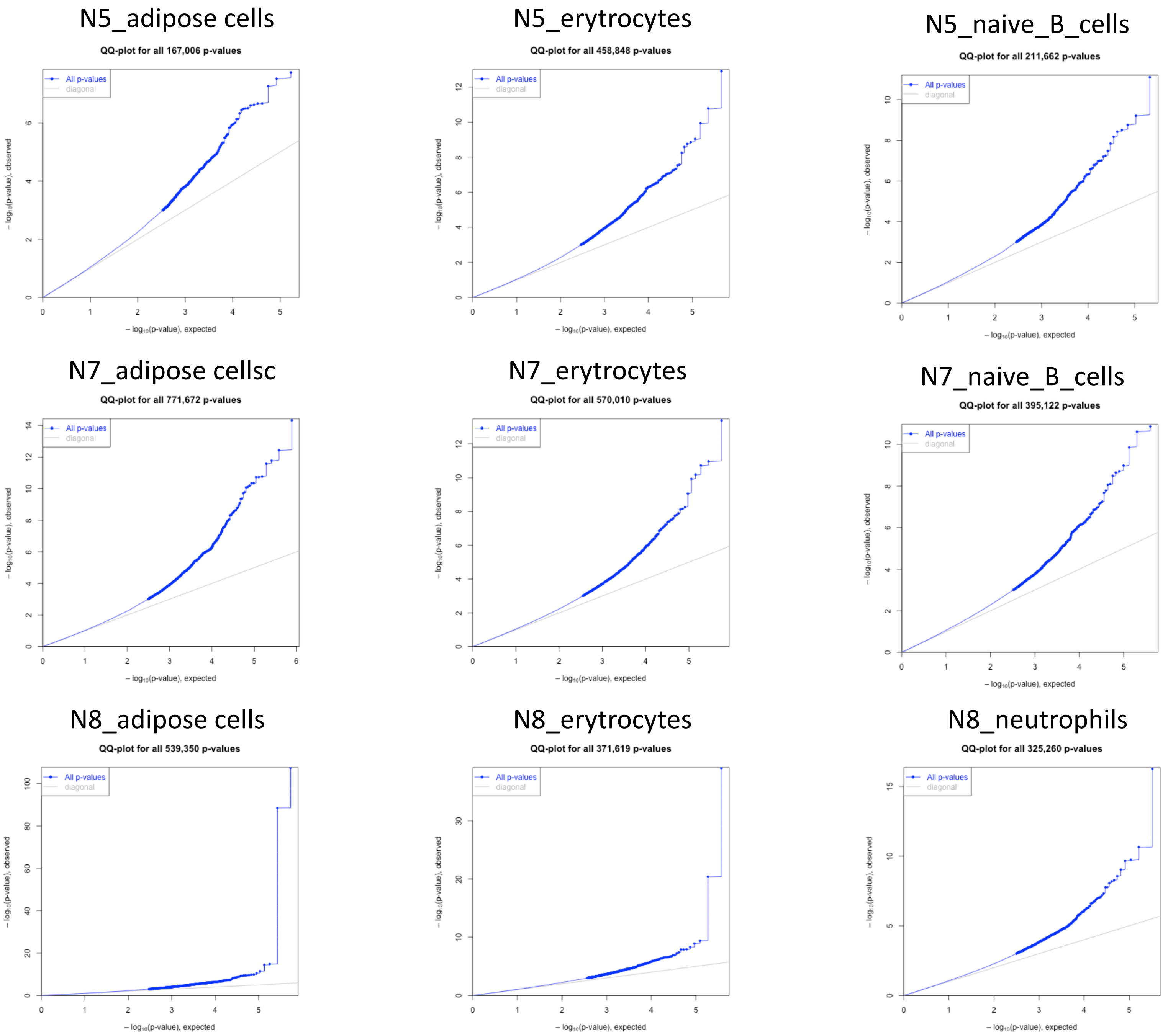


**Supplementary Figure 1.** QQ-plots of the scReQTL P-values.
