## Supplementary Figure 2 for "scReQTL: an approach to correlate SNVs to gene expression from individual scRNA-seq datasets"

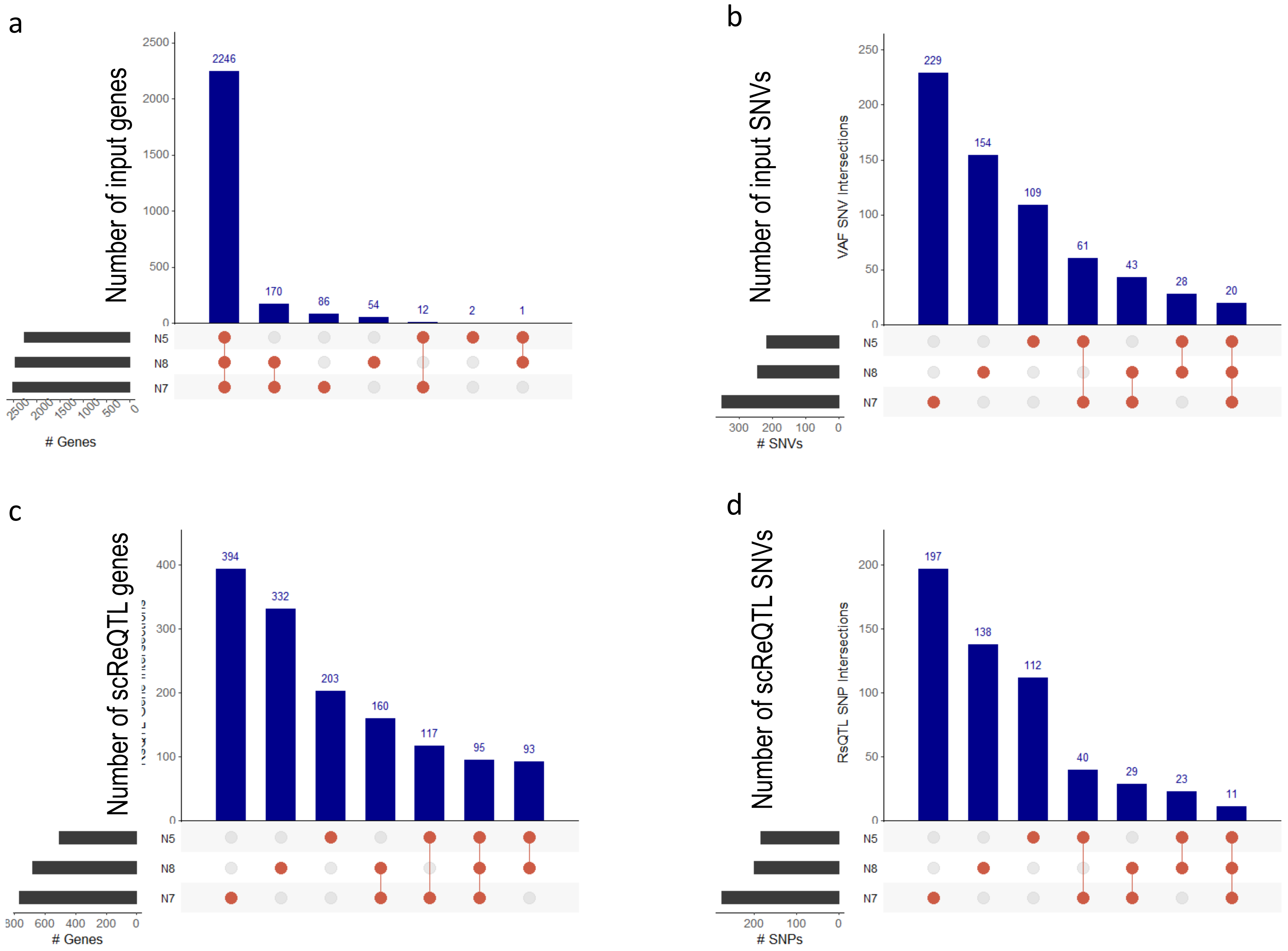


**Supplementary Figure 2**. Relative representation of donor-specific and shared ScReQTL genes and SNVs. **a**) Input genes for scReQTL analysis were shared between the three donors to a large extend. **b**) Input SNV sites for scReQTL analysis were largely donor-specific, with only 20 SNVs shared across the 3 donors. **c**) Shared and donor-specific genes participating in significant scReQTLs. **d**) Shared and donor-specific SNVs participating in significant scReQTLs; 11 of the 20 shared input SNVs participated in scReQTLs shared across the 3 donors.
