## Supplementary Figure 3 for "scReQTL: an approach to correlate SNVs to gene expression from individual scRNA-seq datasets"

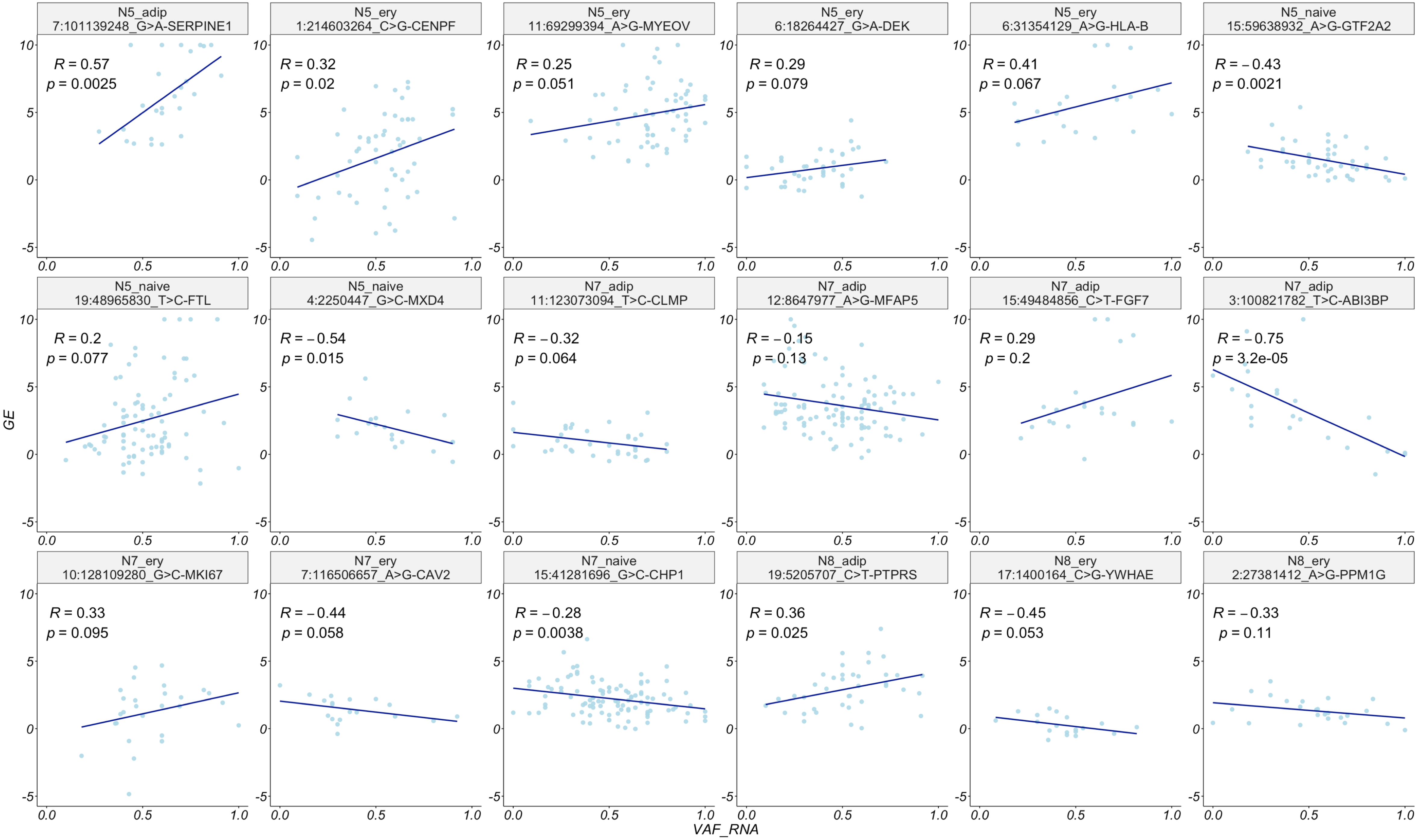


**Supplementary Figure 3**. Examples of scReQTL correlations between SNVs and their harboring genes (cis-scReQTLs) at FDR=0.1. Note that the displayed P-values are calculated based on the input for the plots generated using the R-package ggplot2 and do not represent the FDR—corrected values from the scReQTL analysis performed with Matrix eQTL.
