## Supplementary Figure 4 for "scReQTL: an approach to correlate SNVs to gene expression from individual scRNA-seq datasets"

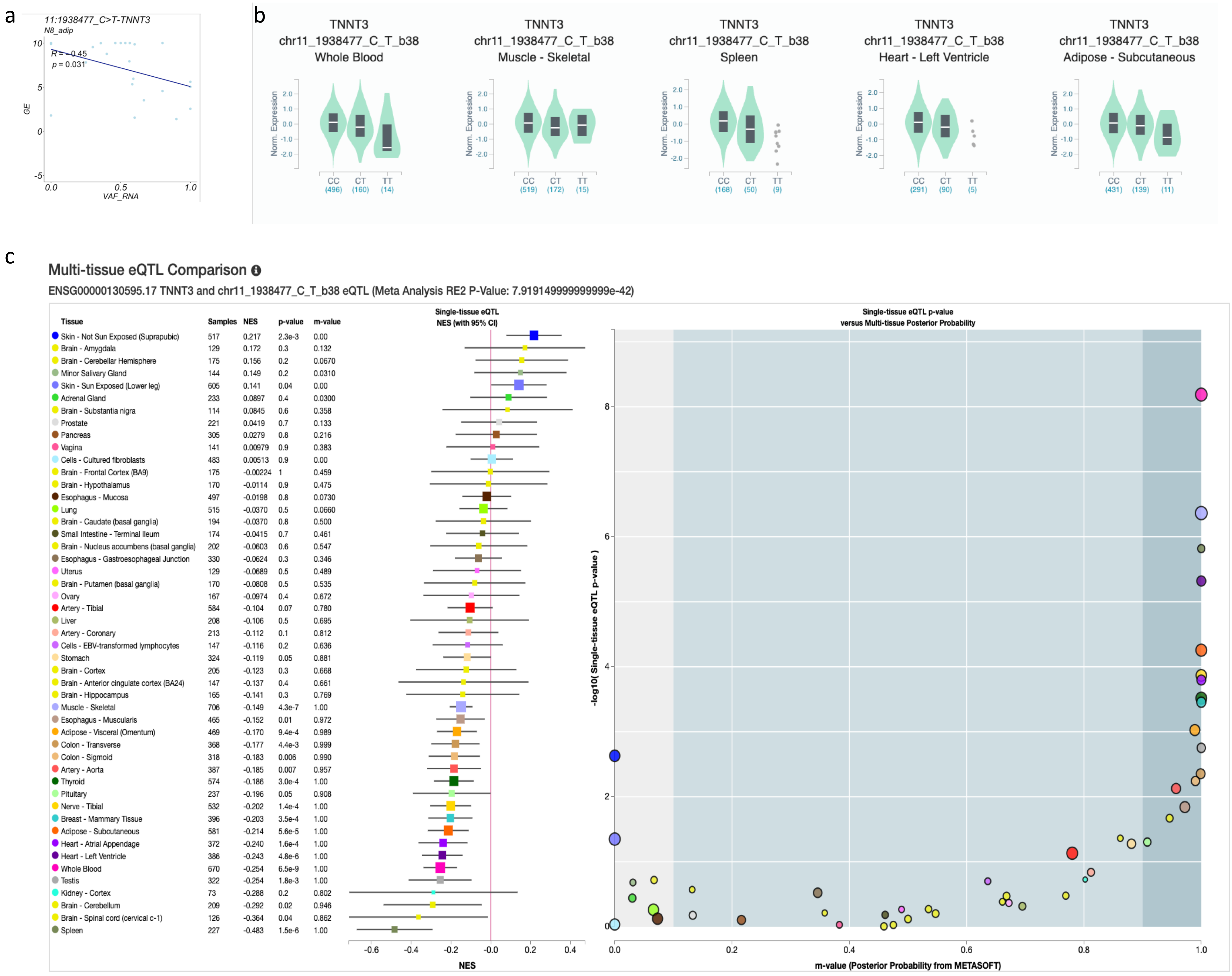


**Supplementary Figure 4**. scReQTL and eQTLs between the SNV at 11:1938477_C>T and its harboring gene *TNNT3* (cis-scReQTL). **a**) scReQTL between the SNV at 11:1938477_C>T and *TNNT3*. **b**) eQTLs between the SNV at 11:1938477_C>T and *TNNT3* reported in the GTEX database in 5 different tissues. In all 5 tissues, the eQTLs were consistent in terms of directionality (negative). c) Multi-tissue comparisons of the eQTL 11:1938477_C>T - *TNNT3*.
