## Supplementary Figure 5 for "scReQTL: an approach to correlate SNVs to gene expression from individual scRNA-seq datasets"

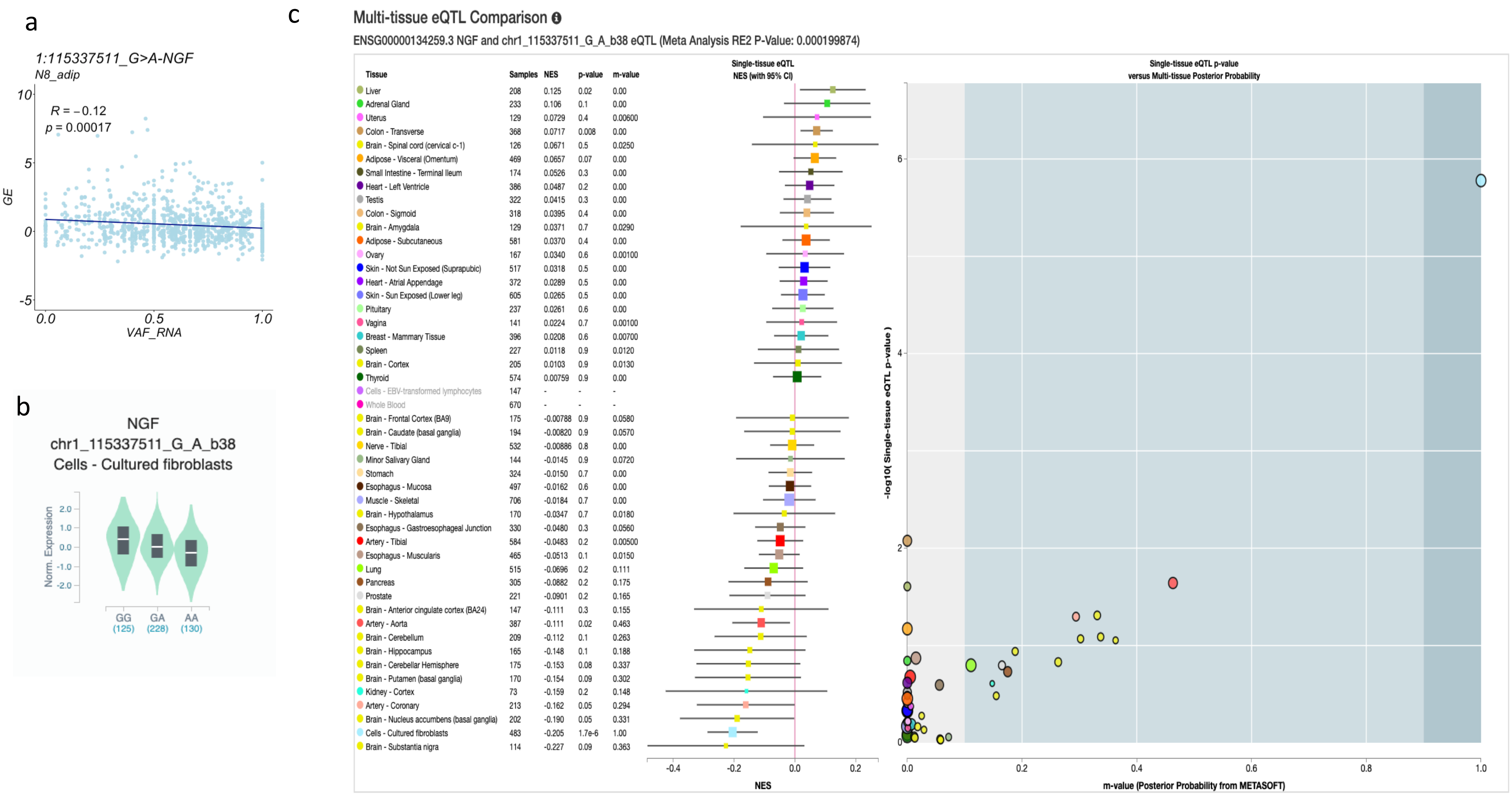


**Supplementary Figure 5**. scReQTL and eQTLs between the SNV at 1:115337511_G>A and the gene *NGF* (trans-scReQTL). **a**) scReQTL between the SNV at 1:115337511_G>A and the gene *NGF*. **b**) eQTL between the SNV at 1:115337511_G>A and the gene *NGF* reported in the GTEX in cultured fibroblasts; the scReQTL and the eQTL were consistent in terms of directionality (negative). **c**) Multi-tissue comparisons of the eQTL 1:115337511_G>A - *NGF*.
