## Supplementary Figure 6 for "scReQTL: an approach to correlate SNVs to gene expression from individual scRNA-seq datasets"

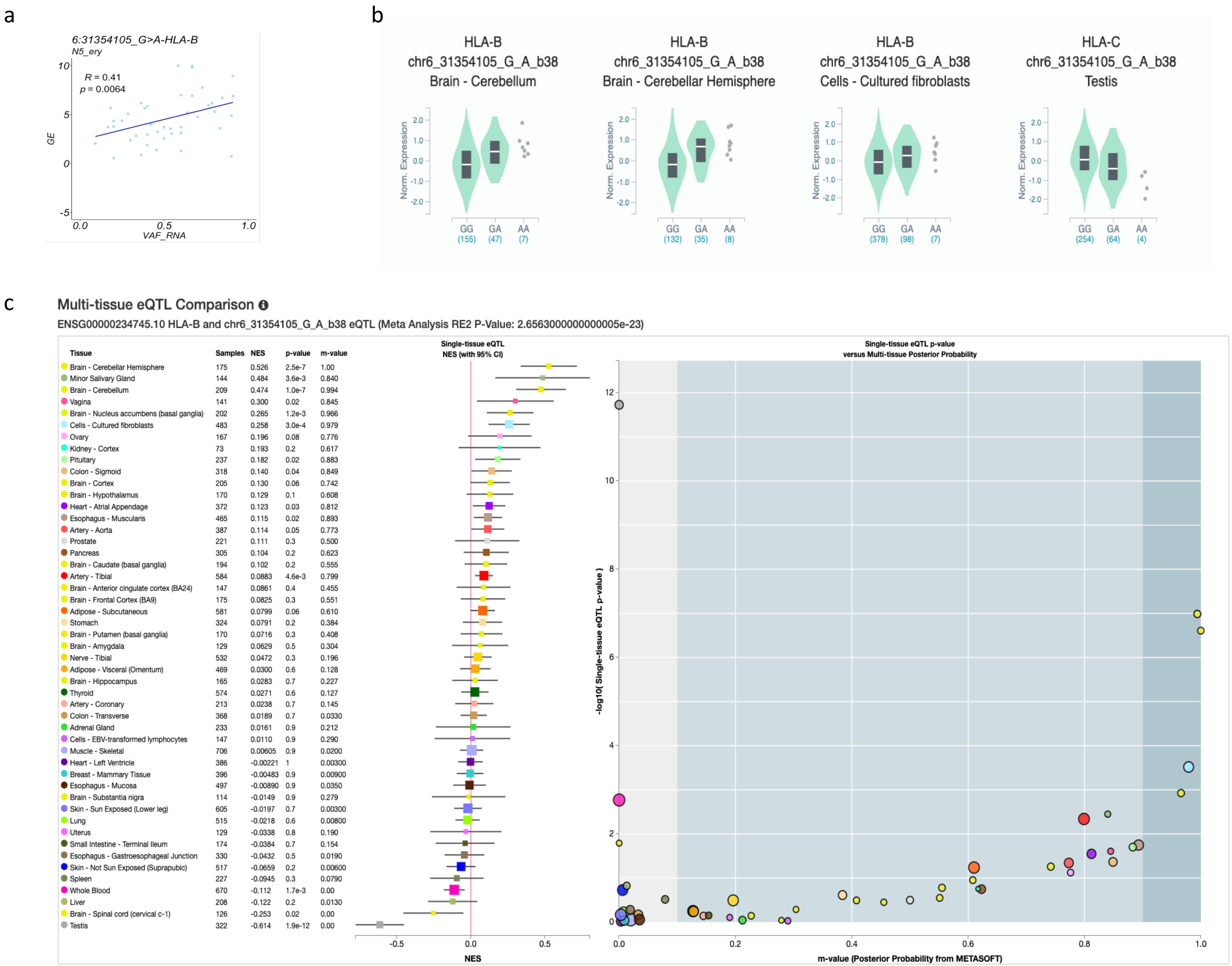


**Supplementary Figure 6**. scReQTL and eQTLs between the SNV at 6:31354105_G>A and its harboring gene *HLA-B* (cis-scReQTL). **a**) scReQTL between the SNV at 6:31354105_G>A and *HLA-B*. **b**) eQTLs between the SNV 6:31354105_G>A and *HLA-B* reported in the GTEX in 4 tissues; the eQTLs correlations were positive in three of the tissues (and agreed with the scReQTL), but in the opposite direction with the eQTL is 1 tissue (testis). **c**) Multi-tissue comparisons of the eQTL at 6:31354105_G>A - *HLA-B*.
