## Supplementary Figure 7 for "scReQTL: an approach to correlate SNVs to gene expression from individual scRNA-seq datasets"

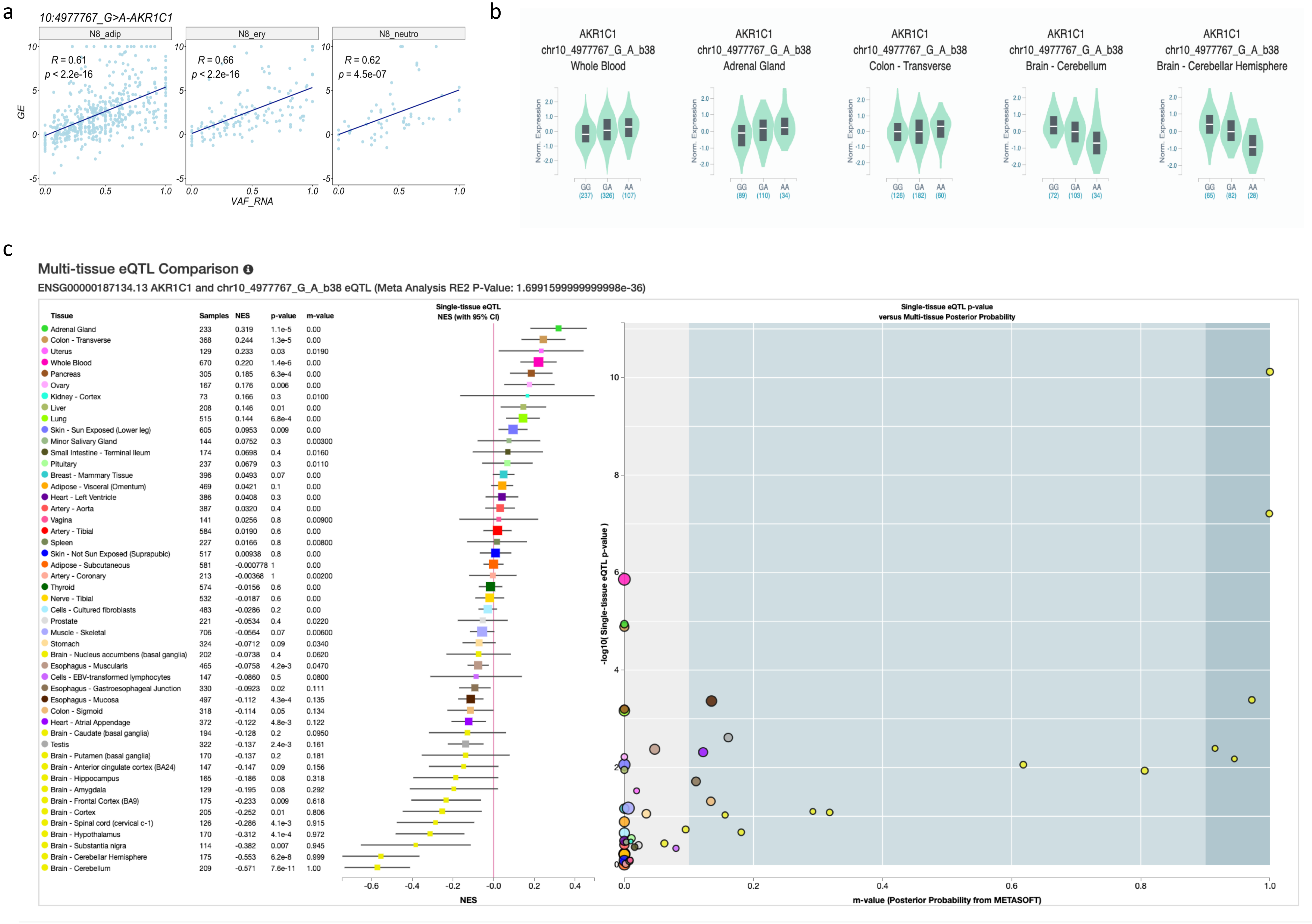


**Supplementary Figure 7**. scReQTL and eQTLs between the SNV at 10:4977767_G>A and its harboring gene *AKR1C1* (cis-scReQTL). **a**) scReQTL between the SNV at 10:4977767_G>A and *AKR1C1*. **b**) eQTLs between the SNV 10:4977767_G>A and *AKR1C1* reported in the GTEX in 5 tissues; the eQTLs correlations were positive in three of the tissues (and agreed with the scReQTL), but in the opposite direction with the eQTL is 2 tissues. **c**) Multi-tissue comparisons of the eQTL at 10:4977767_G>A and *AKR1C1*.
