## Supplementary Figure 8 for "scReQTL: an approach to correlate SNVs to gene expression from individual scRNA-seq datasets"

a

b

c

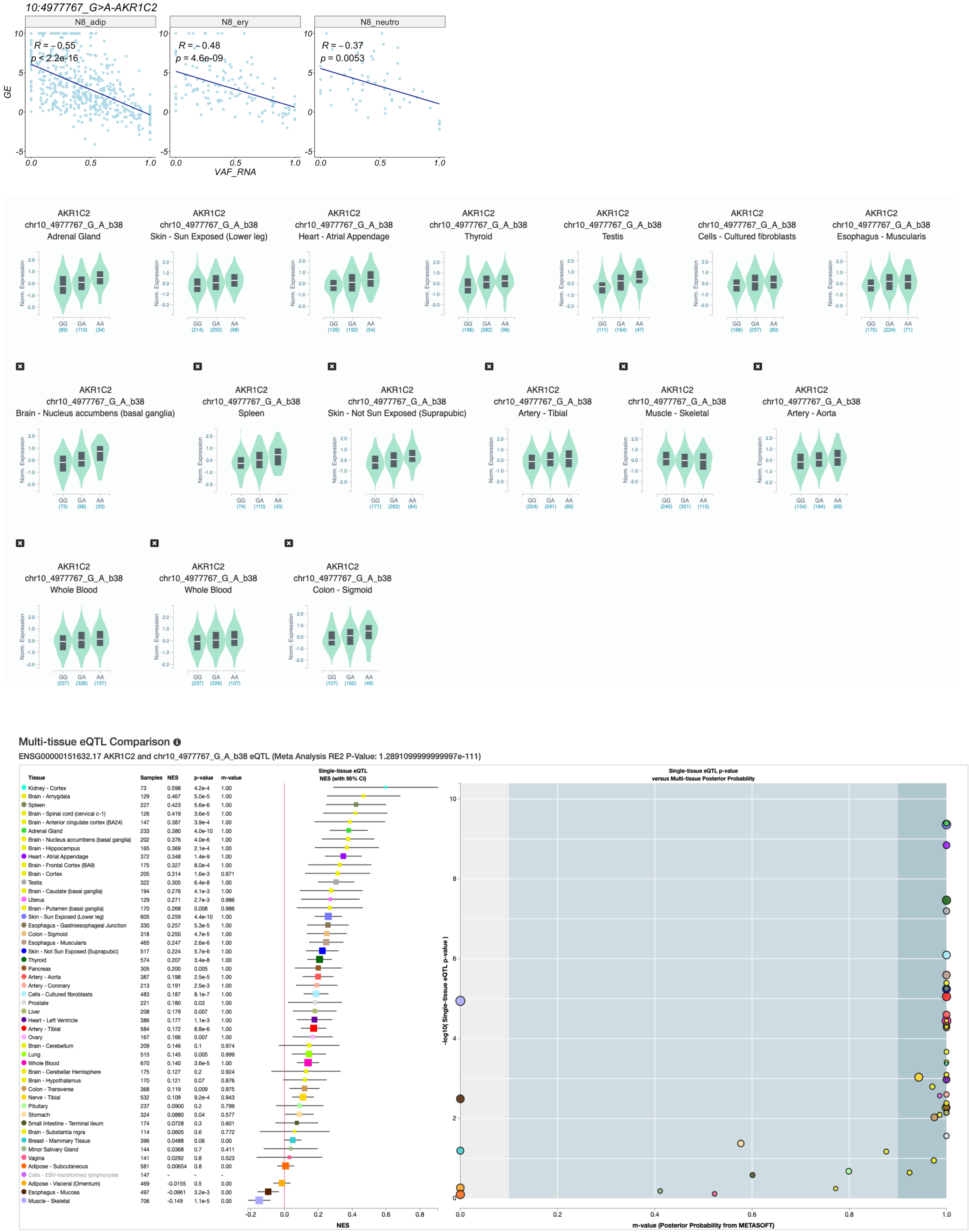


**Supplementary Figure 8.** scReQTL and eQTLs between the SNV at 10:4977767_G>A and *AKR1C2* (trans-scReQTL). **a**) scReQTL between the SNV at 10:4977767_G>A and *AKR1C2*. **b**) eQTLs between the SNV 10:4977767_G>A and *AKR1C2* reported in the GTEX in 6 tissues; the eQTLs correlations were positive in 15 of the tissues, but negative in 1 – skeletal muscle – where they agreed with the scReQTL. **c**) Multi-tissue comparisons of the eQTL 10:4977767_G>A - *AKR1C2* .
