## Supplementary Figure 9 for "scReQTL: an approach to correlate SNVs to gene expression from individual scRNA-seq datasets"

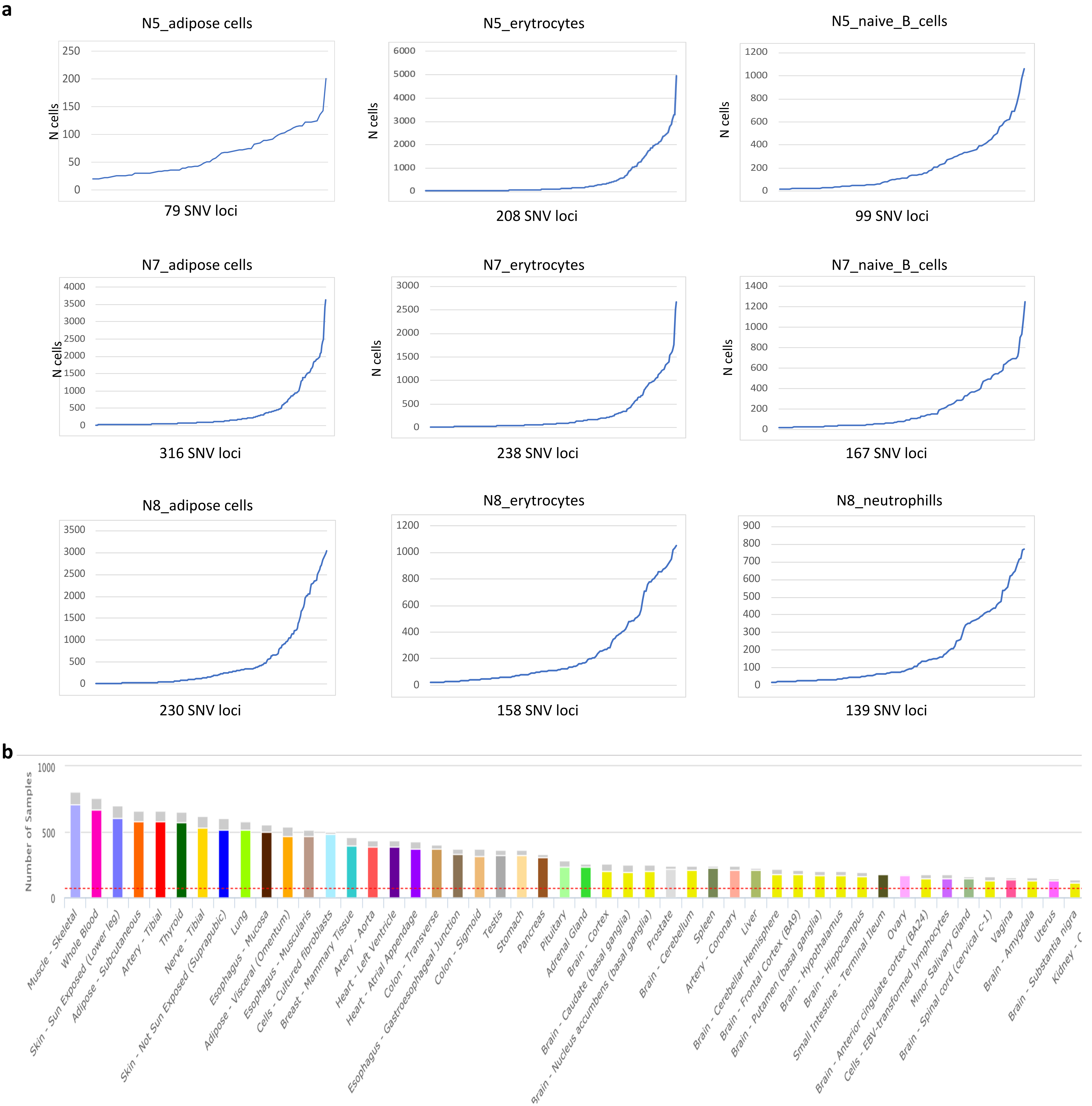


**Supplementary Figure 9**. Number of input cells for scReQTL and input samples (individuals) eQTL in GTEX. **a**) Number of cells with informative VAF_RNA values for scReQTL analysis per donor stratified by cell type. **b**) Number of individuals with genotypes for eQTL analyses in GTEx.
