## Supplementary Figure 10 for "scReQTL: an approach to correlate SNVs to gene expression from individual scRNA-seq datasets"

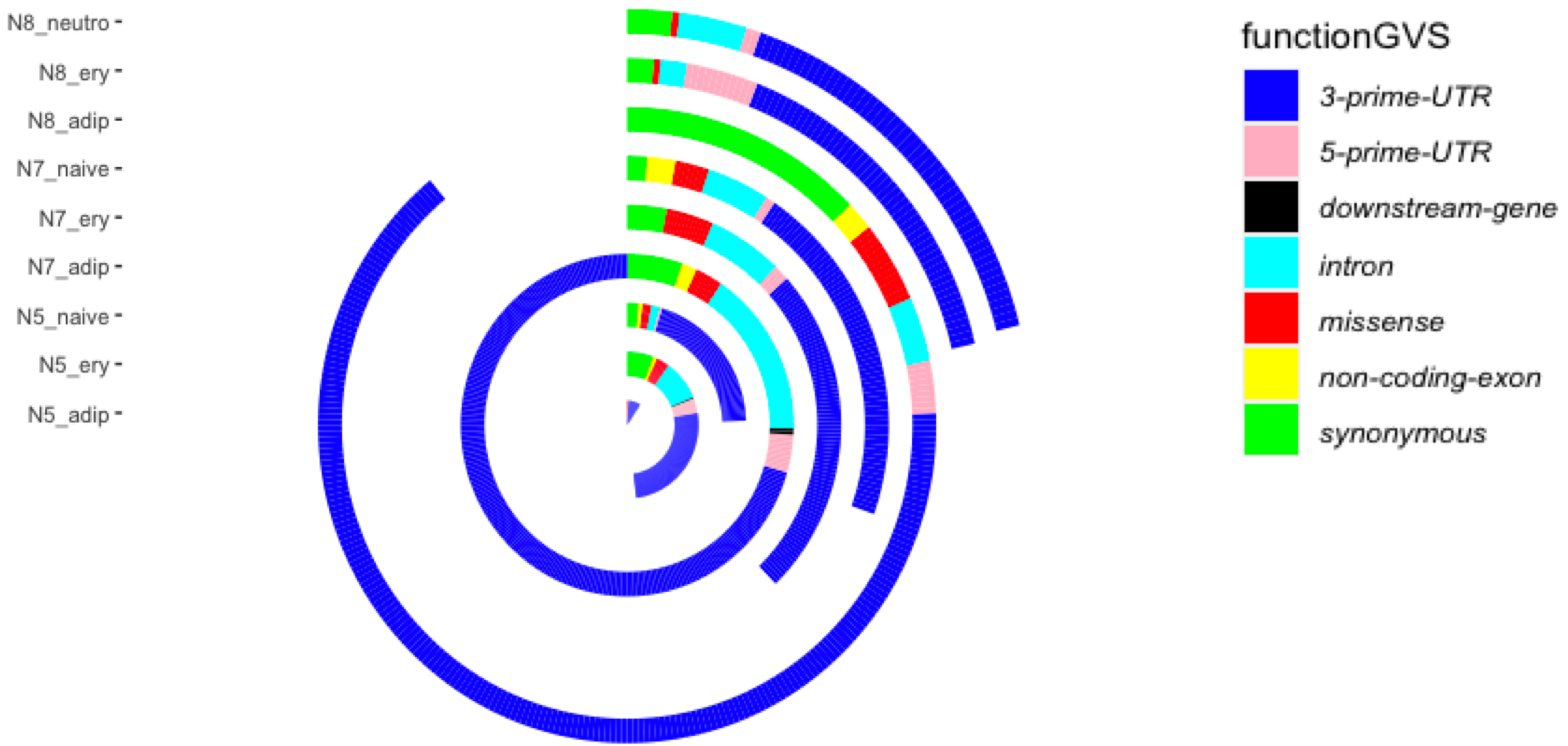


**Supplementary Figure 10**. The majority of the scReQTL SNVs resided in the 3’UTR of their harboring gene (70.2%), followed by exonic SNVs (16.2%), and intronic SNVs (11.2%)
